# Mechanism-based design of DNA-nanoparticle motor with high speed and processivity comparable to motor proteins

**DOI:** 10.1101/2024.05.23.595615

**Authors:** Takanori Harashima, Akihiro Otomo, Ryota Iino

**Affiliations:** Institute for Molecular Science, National Institutes of Natural Sciences, Okazaki, Aichi 444-8787, Japan; Graduate Institute for Advanced Studies, SOKENDAI, Hayama, Kanagawa 240-0193, Japan

**Keywords:** DNA, synthetic molecular motors, DNA nanotechnology, gold-nanoparticles, single-particle tracking

## Abstract

DNA-nanoparticle motor is a burnt-bridge Brownian ratchet moving on RNA-modified surface driven by Ribonuclease H (RNase H), and one of the fastest nanoscale artificial motors. However, its speed is still much lower than those of motor proteins. Here we resolve elementary processes of motion and reveal long pauses caused by slow RNase H binding are the bottleneck. As RNase H concentration ([RNase H]) increases, pause lengths shorten from ∼100 s to ∼0.1 s, while step sizes are constant (∼20 nm). At high [RNase H], speed reaches ∼100 nm s^−1^, however, processivity, run-length, and unidirectionality largely decrease. A geometry-based kinetic simulation reveals switching of bottleneck from RNase H binding to DNA/RNA hybridization at high [RNase H], and trade-off mechanism between speed and other performances. A mechanism-based newly-designed motor with 3.8-times larger DNA/RNA hybridization rate simultaneously achieves 30 nm s^−1^ speed, 200 processivity, and 3 μm run-length comparable to motor proteins.

## Introduction

Engineering artificial molecular motors with performances comparable to motor proteins is an important challenge to understand their operational and design principles and to implement them in biological and abiotic systems^1–5^. DNA is not only the fundamental molecule of living organisms but also a versatile building block used to engineer a variety of artificial motors such as DNA walkers and spiders^6–8^. These artificial motors have flexibility in design, but exhibit speed of only 0.1 nm s^−1^ at best^6,8–10^, and are much slower than motor proteins that exhibit speed of 10–1000 nm s^−1^ ^11–14^. Furthermore, run-length is limited to the size of complementary DNA/RNA tracks, typically tens to hundreds of nanometers.

Recent efforts have focused on overcoming these drawbacks of DNA-based artificial motors. Choi’s group used RNA-modified carbon nanotube as a track for DNAzyme-based motor, and achieved run-length over 3 µm with the maximum speed of 0.1 nm s^−1^ ^10,15^. Salaita’s group developed a rolling motor which consists of DNA-modified microparticle, RNA-modified substrate, and Ribonuclease H (RNase H)^16,17^. In this motor, DNAs on particle and complementary RNAs on planar substrate surface form multivalent DNA/RNA hybridizations. Then, the motion is initiated by addition of RNase H which hydrolyzes RNA in the DNA/RNA duplex, and multiple reaction cycles occur in parallel at the particle/substrate interface. The DNA-microparticle motor cannot go back to the RNA cleaved sites, shows self-avoiding super-diffusion, and operates as a burnt-bride Brownian ratchet (BBR). The motor achieved speed of 30 nm s^−1^ and run-length of 300 μm with the help of gravity on the micron-sized particle^16^.

Salaita’s group also developed DNA-modified gold nanoparticle (AuNP) motors^18^. The nanoscale motors have a potential to achieve higher speed than micron-sized motors due to fast diffusion^19,20^, although its small mass can result in stochastic detachment from the substrate surface. However, in the previous study of a 50-nm DNA-AuNP motor, the average speed was only 3 nm s^−1^ and lower than those of DNA-microparticle motors and nanoscale motor proteins, although transient 50 nm s^-1^ bursts were observed^18^. This result suggests that one or multiple elementary processes become the bottleneck of motion of the DNA-AuNP motor.

Motor proteins simultaneously realize the high speed and long run-length (high processivity). For example, ATP-driven kinesin-1 moves unidirectionally along microtubule with maximum speed of ∼700 nm s^−1^ and run-length longer than 1 μm (processivity higher than 125)^12,21^. In addition to the fast diffusion, high speed motion of kinesin-1 relies on fast chemical processes including ATP binding, phosphate bond cleavage, and product release with millisecond time scales ^21^, and long run-length (high processivity) relies on the highly-coordinated chemo-mechanical coupling where two motor domains control the timings of chemical and mechanical processes^22^. For the motor proteins, the chemo-mechanical coupling mechanisms have been elucidated by single-molecule techniques which resolve the elementary process of the motion^21,23,24^.

In this study, we applied high-speed/high-precision single-particle tracking and resolved the pauses and steps in the motion of a 100-nm DNA-AuNP motor. We found that the bottleneck of motion is the long pauses caused by slow RNase H binding at nanomolar RNase H concentration ([RNase H]) used in the previous study^18^. As [RNase H] increased, the pause lengths shortened from ∼100 to ∼0.1 s, while the step sizes (∼20 nm) were constant. At micromolar [RNase H], the speed reached ∼100 nm s^−1^, however, processivity, run-length, and unidirectionality largely decreased.

To understand the mechanisms of motion and trade-off between speed and other performances, we also developed a geometry-based kinetic simulation which reproduces experiment quantitatively. The simulation revealed that pauses are caused by multiple DNA/RNA hybridizations between DNA-AuNP and RNA-substrate and steps occur when the number of hybridizations decreases by the progress of RNA hydrolysis. The simulation also revealed that as [RNase H] increases, the bottleneck switches from RNase H binding to DNA/RNA hybridization, resulting in low processivity, short run-length, and low unidirectionality. Furthermore, the simulation predicted that large DNA/RNA hybridization rate improves the trade-off.

Finally, based on the prediction of the simulation, we designed new DNA-AuNP motor by using a DNA/RNA sequence which shows 3.8-times larger hybridization rate than the original one. As a result, the newly-designed motor simultaneously achieved speed of 30 nm s^−1^, processivity of 200, and run-length of 3 μm comparable to motor proteins. Our study demonstrates a mechanism-based strategy to improve the performance of artificial molecular motors.

## Results

### Stepping motion of DNA-AuNP motor visualized by high-speed/high-precision tracking

The 100-nm DNA-AuNP motor was prepared by using the DNA/RNA sequences similar to the previous study (Supplementary Table 1)^18^. The densities of the DNA on AuNP surface and the RNA on glass substrate surface were 0.11 ± 0.02 molecules nm^−2^ and 0.022 ± 0.008 molecules nm^−2^, respectively (Supplementary Fig. 1 and 2).

For high-speed/high-precision single-particle tracking, we obtained image sequences at 20–1000 frames per second (fps), much higher rate than the previous study (0.2–2 fps)^18^, by using a total internal reflection dark-field microscopy (Fig. 1a)^25,26^. For long-term measurements at 36 nM RNase H, similar [RNase H] used in the previous study^18^, the localization precision was 3.1 nm. At other [RNase H]s (see below), the localization precision was 0.8–1.0 nm (Supplementary Fig. 3).

**Figure 1.**
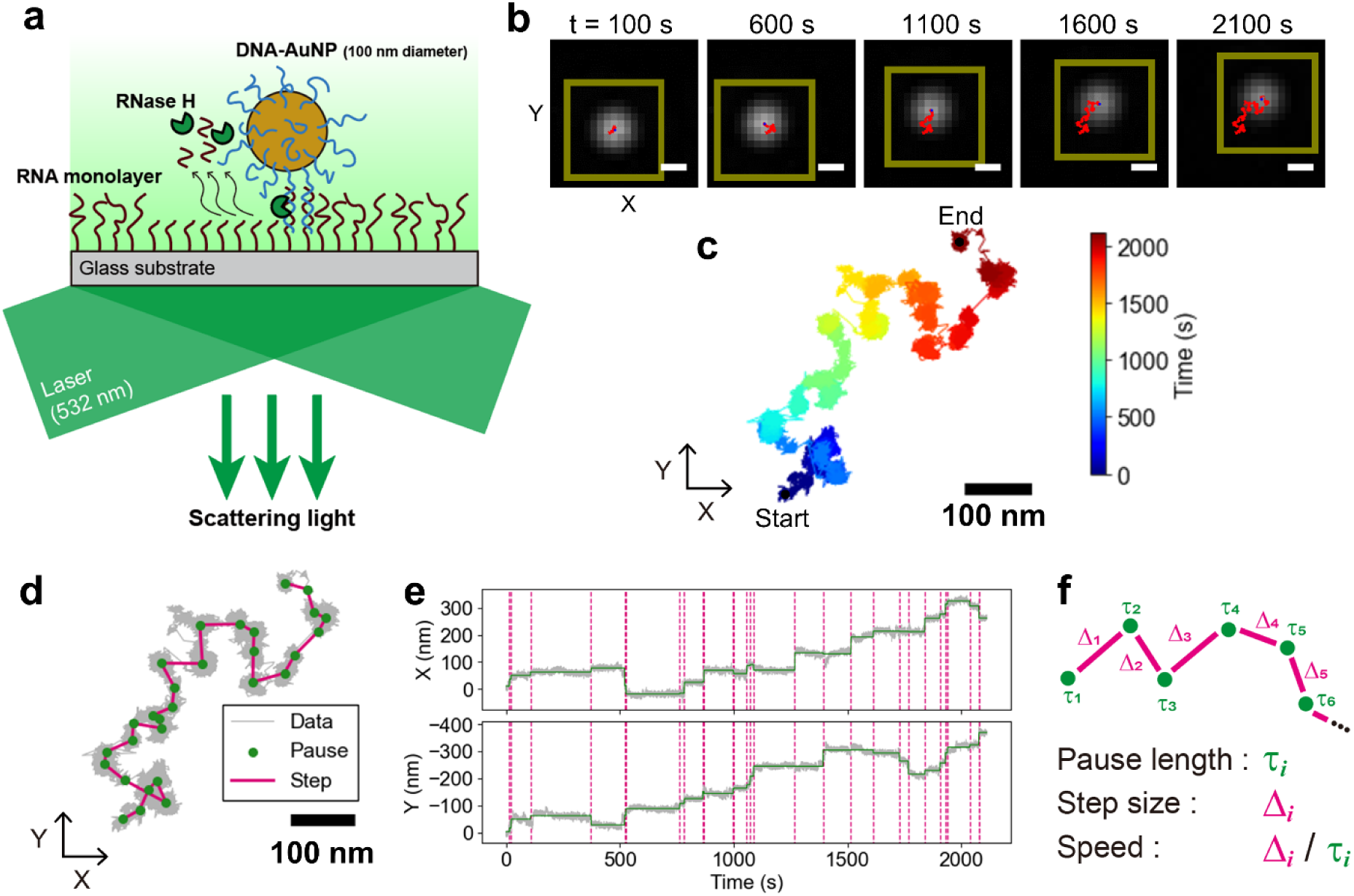
Observation of elementary processes of motion of DNA-AuNP motor. **(a)** Schematic illustration of the experimental system for high-speed/high-precision single particle tracking of DNA-AuNP motor. DNAs on the AuNP surface are hybridized with RNAs on the glass surface. RNAs that hybridized with DNAs are selectively hydrolyzed by RNase H. The motion of DNA-AuNP motor is visualized by using total-internal reflection dark-field microscopy. **(b)** An image sequence of motion of DNA-AuNP motor on RNA surface at 36 nM RNase H. Recording rate: 20 fps. Red lines indicate trajectories of the particle centroid. Dark yellow squares show the regions of interest to calculate the centroid for each frame. Scale bar: 200 nm. **(c)** Whole trajectory of the centroid of DNA-AuNP motor shown in (b). **(d)** Detected pauses (green dots) and steps (magenta lines) superimposed to raw trajectory shown in (c). **(e)** Time-course of X- and Y-coordinates. Green solid lines are fittings to the raw trajectories, and magenta dashed lines indicate the positions of detected steps. **(f)** Schematic illustration and definitions of pause length, step size, and speed.

Without RNase H, the motors were stationary on the RNA substrate, and exhibited 2D motions immediately after the RNase H addition (Fig. 1b, Supplementary Video 1). During observations, most motors kept moving and some detached from the substrate (the ratio of detachment depended on [RNase H], see below). Interestingly, 2D trajectories of the motor clearly showed pauses and steps (Fig. 1c). These stepping motions were resolved for the first time.

Then, the pauses and steps were detected objectively by using a step-finding algorithm (Fig. 1d and e)^27^. Individual pause lengths and step sizes were collected for subsequent statistical analysis. We also calculated the speed by dividing each step size with each pause length (Fig. 1f). At 36 nM RNase H, the median pause length, step size, and speed were 70 s, 24 nm, and 0.33 nm s^−1^, respectively (Fig. 2a-c). Therefore, we conclude that low speed is caused by the long pauses.

**Figure 2.**
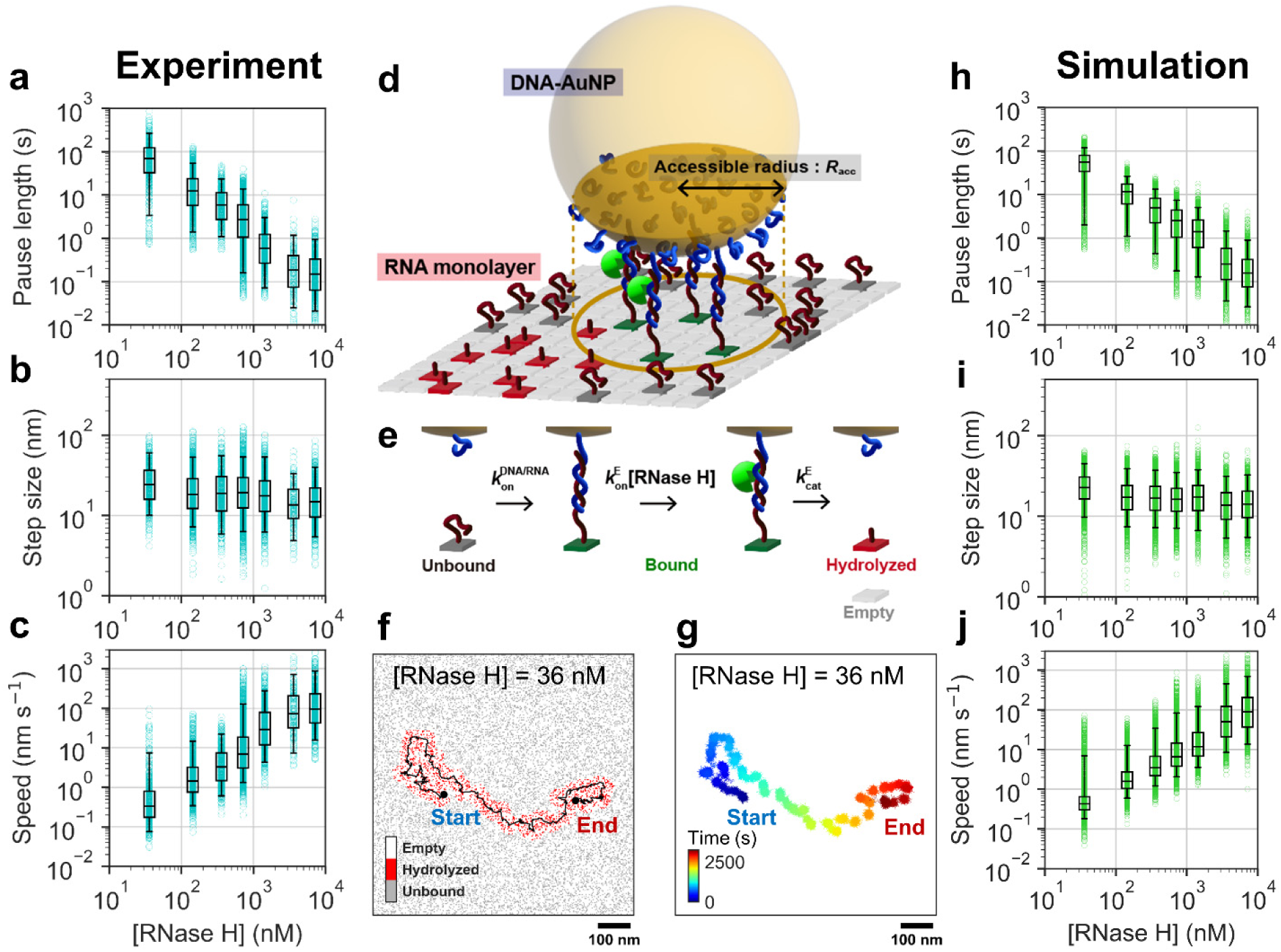
[RNase H] dependence of pause length, step size, and speed measured by single-particle tracking experiments and geometry-based kinetic simulations. **(a-c)** [RNase H] dependence of pause length (a), step size (b), and speed (c) in the experiments. Individual data are shown by cyan circles. Box plot shows interquartile range (IQR). Error bars show 5^th^/95^th^ percentile for each [RNase H] condition. Recording rates were 20, 20, 250, 250, 250, 1000, 1000 fps for 36, 144, 360, 720, 1440, 3600, and 7200 nM RNase H, respectively. **(d)** Schematic illustration of geometry-based kinetic simulation. Hybridization between RNA and DNA occurs stochastically within an area with the accessible radius (*R*_acc_) of 28.0 nm, where single DNA-AuNP motor is accessible to the RNA surface (dark yellow circle). Multivalent DNA/RNA hybridizations immobilize the motor and result in the pauses, and the steps occur after the successive degradations of RNAs by RNase H. **(e)** Kinetic model at each RNA-modified site on the surface. *k*_on_^DNA/RNA^, *k*_on_ ^E^, and *k*_cat_^E^ correspond to the rate constants of DNA/RNA hybridization, RNase H binding to DNA/RNA duplex, and RNA hydrolysis, respectively. Empty sites are inert in the simulation. The values of *k*_on_^DNA/RNA^, *k*_on_ ^E^, and *k*_cat_^E^ were set to 0.3 s^−1^, 1.0 x 10^6^ M^-1^s^-1^ and 4 s^−1^, respectively. **(f)** Representative trajectory of the DNA-AuNP motor translocation superimposed on spatial distribution of RNA states at 36 nM RNase H. **(g)** Whole trajectory of the centroid after the addition of the Gaussian noise (*σ*_noise_ = 6 nm). **(h-j)** [RNase H] dependence of pause length, step size, and speed in the simulation. Individual data are shown by green circles.

### Speed increases as [RNase H] increases and reaches 100 nm s^−1^

To gain information on the chemical processes which lengthen the pauses, we next conducted fluorescence bulk assays of DNA/RNA hybridization and RNA hydrolysis (Extended Data Fig. 1a, b). We found that apparent hybridization rate increased linearly up to 100 nM DNA (RNA concentration was fixed at 10 nM) (Extended Data Fig. 1c, d). Estimated rate constants for DNA/RNA hybridization (*k*_on_ ^DNA/RNA^) and dissociation (*k*_off_ ^DNA/RNA^) were (3.9 ± 0.5) **×** 10^5^ M^−1^s^−1^ and (2.0 ± 0.9) **×** 10^−3^ s^−1^, respectively (Supplementary Table 2). On the other hand, apparent RNA hydrolysis rate showed a saturation above micromolar RNase H (Extended Data Fig. 1e, f). Rate constants for RNase H binding (*k*_on_ ^E^) and RNA hydrolysis (*k*_cat_^E^) were estimated to be (2.2 ± 0.3) **×** 10^6^ M^−1^s^−1^ and 1.6 ± 0.1 s^−1^, respectively, indicating that RNase H binding is the bottleneck of RNA hydrolysis at nanomolar [RNase H].

We then conducted single-particle tracking at different [RNase H]s from 36 to 7200 nM (Supplementary Fig. 4, Supplementary Video 1, 2, 3). Resulting pause lengths, step sizes, and speeds are summarized (Fig. 2a-c, Supplementary Fig. 5). We found that the pause length largely shortened from ∼100 s to ∼0.1 s as [RNase H] increased (Fig. 2a), while the step size (∼20 nm) did not change largely (Fig. 2b). As results, the speed increased as [RNase H] increased (Fig. 2c). Remarkably, at highest 7200 nM RNase H, the speed reached 100 nm s^−1^.

### Geometry-based kinetic simulation reproduces stepping motion and [RNase H] dependence

To understand the mechanism of stepping motion of the DNA-AuNP motor, we developed a geometry-based kinetic simulation (Supplementary Note, Supplementary Fig. 6, Supplementary Table 3). Briefly, individual DNAs on the AuNP faced with the RNA-substrate are represented by the square grids in the accessible region (Fig. 2d, Supplementary Fig. 7), and RNAs are randomly distributed on the same size grids on the substrate surface to reproduce density difference between DNA and RNA. Then, reactions are stochastically progressed in parallel at individual RNA sites in the accessible region. The reaction model is composed of three sequential processes: 1) DNA/RNA hybridization (*k*_on_^DNA/RNA^), 2) RNase H binding to DNA/RNA duplex (*k*_on_^E^[RNase H]), and 3) RNA hydrolysis (*k*_cat_^E^) (Fig. 2e). We do not consider dissociation of the intact DNA/RNA duplex^28^, because *k*_off_ ^DNA/RNA^ estimated in the bulk assay was very small (Supplementary Table 2). The XY-coordinates of the DNA-AuNP are randomly varied within the overlapped region among the mobile regions of all RNA sites bound with DNA (Supplementary Fig. 8).

As a result, simulated trajectories showed stepping motions similar to the experiments (Fig. 2f, Supplementary Video 4, 5, 6). Note that in the simulation, the raw trajectories showed much smaller fluctuations than the experiments during the pauses (Fig. 2f), because the motor is completely stationary if there is no overlap among the mobile regions of all bound sites. Thus, in order to reproduce the experiments more precisely, we added Gaussian noise to the raw simulation trajectories (Fig. 2g, Supplementary Note, Supplementary Fig. 9 and 10). Furthermore, based on the values obtained by the bulk assays (Supplementary Table 2), we optimized kinetic parameters used in the simulation by using a simulation-based fitting (Supplementary Fig. 11-16). In short, small numbers of the simulations were conducted with different kinetic parameters, and the mean square errors (MSE) of pause length and step size between the simulation and experiment were calculated. The optimal set of the kinetic parameters which showed the minimum MSE was *k*_on_ ^DNA/RNA^ = 0.3 s^−1^, *k*_off_ ^E^ = 1.0 × 10^6^ M^−1^s^−1^, and *k*_cat_^E^ = 4.0 s^−1^ (Supplementary Fig. 16, Supplementary Table 3). Then, numbers of simulations were increased and each trajectory was analyzed again. As results, the pause lengths, step sizes, and speeds obtained by the simulation showed excellent agreement with those obtained by the experiment at all [RNase H]s (Fig. 2h-j).

### [RNase H]-dependent coupling between pause-step cycle and elementary chemical processes

We investigated how the coupling between pause-step cycle and elementary chemical processes depends on [RNase H]. In our simulation, *k*_on_^E^[RNase H] increases as [RNase H] increases, while *k_on_* ^DNA/RNA^ and *k*_cat_^E^ are constant (Supplementary Table 3). Therefore, relative amplitude among these rates varies depending on the [RNase H] (Fig. 3a, regions (I)–(III)). The boundaries between (I) and (II), and (II) and (III) are 300 and 4000 nM RNase H, respectively. We then analyzed time course of the numbers of RNAs unbound (*N*_unbound_) and bound (*N*_bound_) with DNAs and hydrolyzed RNAs (*N*_hydrolyzed_) in the accessible region at 36, 720, and 7200 nM RNase H, each of which corresponds to regions (I), (II), and (III), respectively (Fig. 3b, Supplementary Video 4, 5, 6). We focused on time points corresponding to (i) just after the step, (ii) during the pause, (iii) just before the next step, and (iv) just after the next step (= (i)). The spatial distributions of RNA states at each time point (i)–(iv) are also shown.

**Figure 3.**
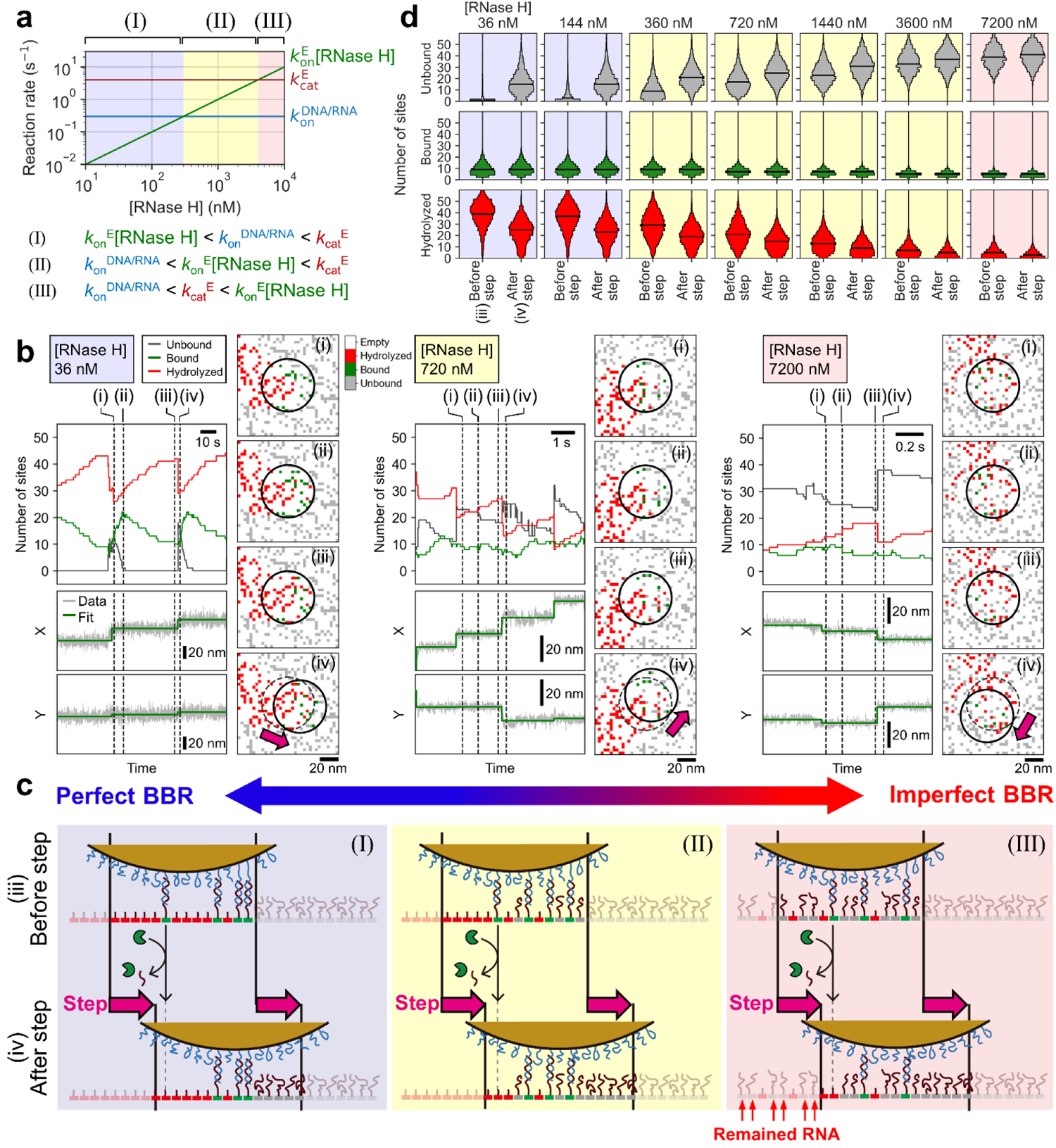
Stepping mechanisms depending on [RNase H]. **(a)** Relationships among three reaction rates considered in this study *k*onDNA/RNA, *k*onE [RNase H], and *k*_cat_^E^) depending on [RNase H]. Blue, yellow, and red shaded regions correspond to different order of magnitude of these rates (I)–(III). **(b)** Time courses of the numbers of unbound RNAs, RNAs bound with DNAs, and hydrolyzed RNAs at 36 (left), 720 (middle), and 7200 (right) nM RNase H. X- and Y-coordinates of the motion are also shown. Black dashed lines labeled by (i)–(iv) show the representative frames at (i, iv) just after step, (ii) during pause, (iii) just before step. Spatial distributions of RNA around DNA-AuNP motor corresponding to the frames of (i)–(iv) are shown in the right of the time courses. Black circles represent accessible region of the DNA-AuNP motor. Gray dashed circles in (iv) show the accessible region in (iii). Pink arrows indicate the step directions. **(c)** Schematic illustrations of the stepping mechanisms at different [RNase H]s at regions (I)–(III). **(d)** Violin plots of the number of RNA sites within the accessible region just before and after steps. Black solid line shows the median value.

At 36 nM RNase H (Fig. 3b, left), at (i), DNAs contacted with new unbound RNAs and started to form new hybridizations. Note that the unbound and hydrolyzed RNAs distributed in the forward and backward half of the accessible region, respectively, because the step size (∼20 nm) is comparable to the radius of accessible region (*R*_acc_ = 28.0 nm, Supplementary Fig. 7). Between (i) and (ii), *N*_bound_ increased faster than *N*_hydrolyzed_, because *k*_on_ ^DNA/RNA^ is much larger than *k*_on_ ^E^[RNase H] at region (I). At (ii), DNA/RNA hybridizations were completed. Between (ii) and (iii), *N*_bound_ and *N*_hydrolyzed_ decreased and increased, respectively. Then, between (iii) and (iv), the step occurred again when *N*_bound_ decreased to less than 10. At (iv), DNAs became accessible to new unbound RNAs again. Importantly, almost all RNAs under the trajectory of DNA-AuNP were hydrolyzed during the stepping motion as the “perfect” BBR (Fig. 3c, left).

At 720 nM RNase H (Fig. 3b, center), the hybridization of unbound RNAs and hydrolysis of bound RNAs proceeded in parallel between (i) and (iii), resulted in nearly constant *N*_bound_ around 10. Interestingly, there were some unbound RNAs in the forward half of the accessible region at (iii). Then, many of them moved to the backward half after the step ((iv)), and hydrolyzed before the next step. Thus, the BBR mechanism still works (Fig. 3c, center).

At 7200 nM RNase H (Fig. 3b, right), on the other hand, the *N*_bound_ was kept lower than 10 throughout (i) to (iv), because *k*_on_ ^DNA/RNA^ is much lower than *k*_on_^E^[RNase H] and *k*_cat_^E^. This resulted in high probability of detachment and low processivity and short run-length (see below). Moreover, there were many unbound RNAs in the entire accessible region even at (iii), and remained intact under the trajectory of DNA-AuNP (Fig. 3c, right). This “imperfect” BBR caused frequent backward steps which resulted in low unidirectionality (see below).

We also performed statistical analysis of the *N*_unbound_, *N*_bound_, and *N*_hydrolyzed_ just before (iii) and after (iv) the steps (Fig. 3d). Notably, at all [RNase H]s, at both (iii) and (iv), the *N*_bound_ was always around less than 10, consistent with the notion that the steps occur when *N*_bound_ <10. On the other hand, at (iii), *N*_unbound_ increased from nearly 0 to 40, and *N*_hydrolyzed_ decreased from 40 to 5 as [RNase H] increased, reflecting the shift from the perfect to imperfect BBR mechanism.

### Trade-off between speed and processivity/run-length and mechanism of detachment

Next, we examined [RNase H] dependence of processivity and run-length (Fig. 4). For experimental data, time courses of the fraction of motors remained attached on RNA substrate were fitted with single exponential decay functions (Fig. 4a) to estimate time constants before detachment (τ_detach_) (Fig. 4b)^18,29^. Processivity and run-length estimated from τ_detach_, mean pause length, and mean step size (Fig. 4c) are shown in Fig. 4d and e, respectively. When [RNase H] < 1000 nM, the motor showed high processivity of 100∼200 and long run-length of 3∼4 μm. However, when [RNase H] > 1000 nM, processivity and run-length largely decreased, and importantly, similar [RNase H] dependence was observed in the simulation (Extended Data Fig. 2a, b).

**Figure 4.**
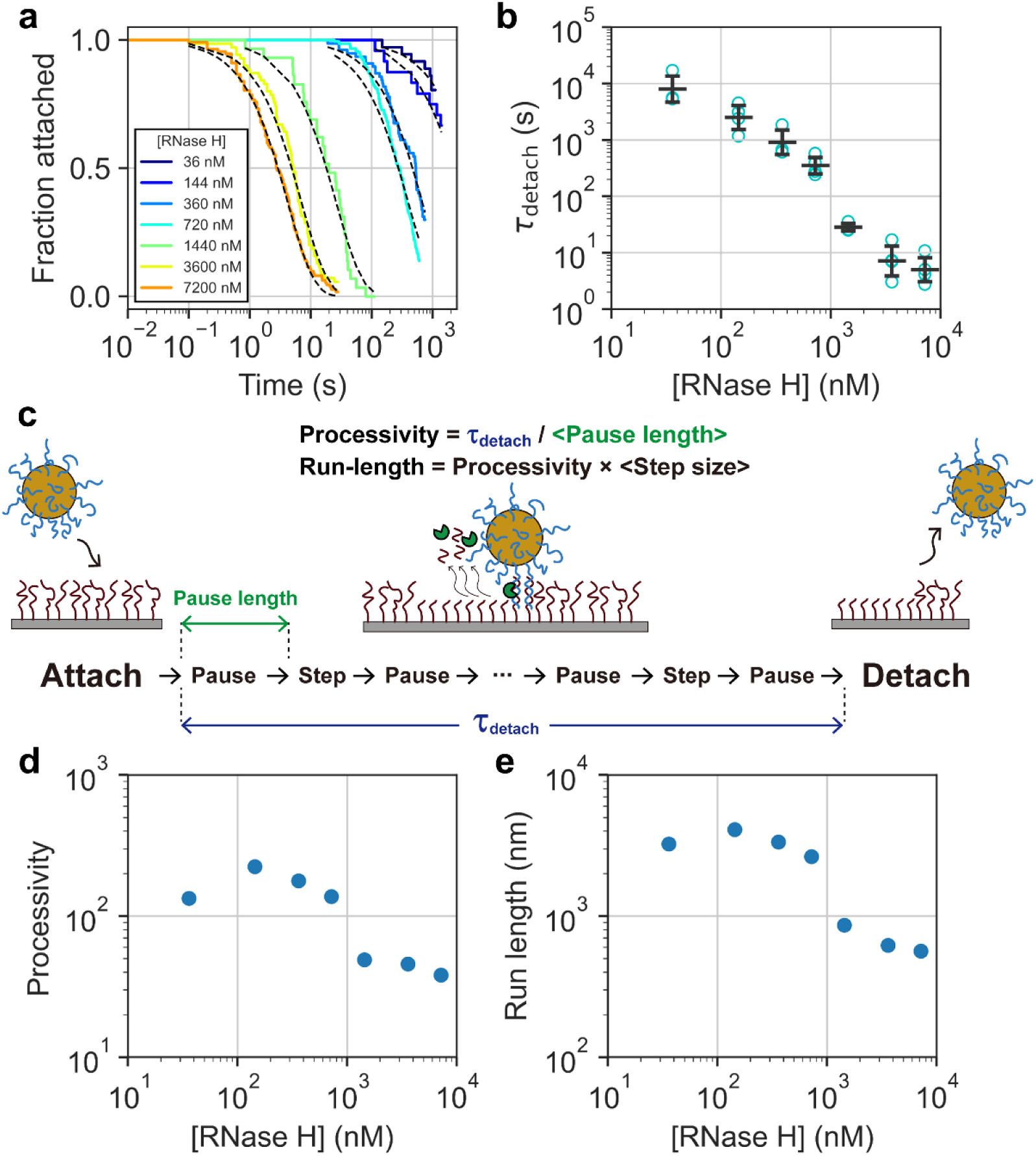
[RNase H] dependence of processivity and run-length of the experimental data. **(a)** Typical plots of fraction of motors remained attached to the RNA surface as a function of time. Each plot was fitted by single exponential decay function, exp(−t/*τ*_detach_), that is superimposed as the black dashed line. **(b)** [RNase H] dependence of *τ*_detach_. Black bars and error bars correspond to mean ± s.d. The mean and s.d. values were calculated from at least three independent experiments for each [RNase H] condition (cyan circles). **(c)** Schematic illustration of whole kinetics from attachment to detachment from RNA surface. Processivity was defined as the number of steps before detachment and calculated by dividing the *τ*_detach_ by mean pause length. Run-length was estimated by multiplying the processivity with mean step size. **(d, e)** [RNase H] dependence of processivity (d) and run-length (e).

Then, we investigated the mechanism of detachment in the simulation. We found that there are two modes of detachment associated with processivity and run-length (Extended Data Fig.2c, d). One is caused by the entrapment of DNA-AuNP into the area where surrounding RNAs are already hydrolyzed (entrap detachment), and another is caused by the stochastic loss of all DNA/RNA hybridizations (stochastic detachment). We also analyzed the ratio (*P*_unbound_) of unbound RNAs to unbound and hydrolyzed RNAs around the DNA-AuNP just before the detachment (Extended Data Fig. 2e, f). We found that as [RNase H] increased, *P*_unbound_ increased in a sigmoidal fashion. When [RNase H] was higher than 1000 nM, *P*_unbound_ was larger than 0.5, indicating that stochastic detachment is major and results in low processivity and short run-length.

### Trade-off between speed and unidirectionality

The DNA-nanoparticle and -microparticle motors show super-diffusion, where the motions are biased forward by suppressing backward motions as 2D BBR^16,18^. In this study, to evaluate super-diffusive motion directly, we analyzed step angles (Δ*θ*) between pauses (Fig. 5a, see Supplementary Note and Supplementary Fig. 17 for the analysis of mean square displacement). For both the experiment and simulation, when [RNase H] < 1000 nM, distribution of the Δ*θ* showed a peak around 0°, indicating step directions are biased forward (Fig. 5b and d). Forward-biased steps are more clearly shown in the polar plots (Fig. 5c and e). When [RNase H] > 1000 nM, on the other hand, the number of backward steps increased and the Δ*θ* distribution became more uniform, indicating low unidirectionality at high [RNase H] (Supplementary Fig. 18).

**Figure 5.**
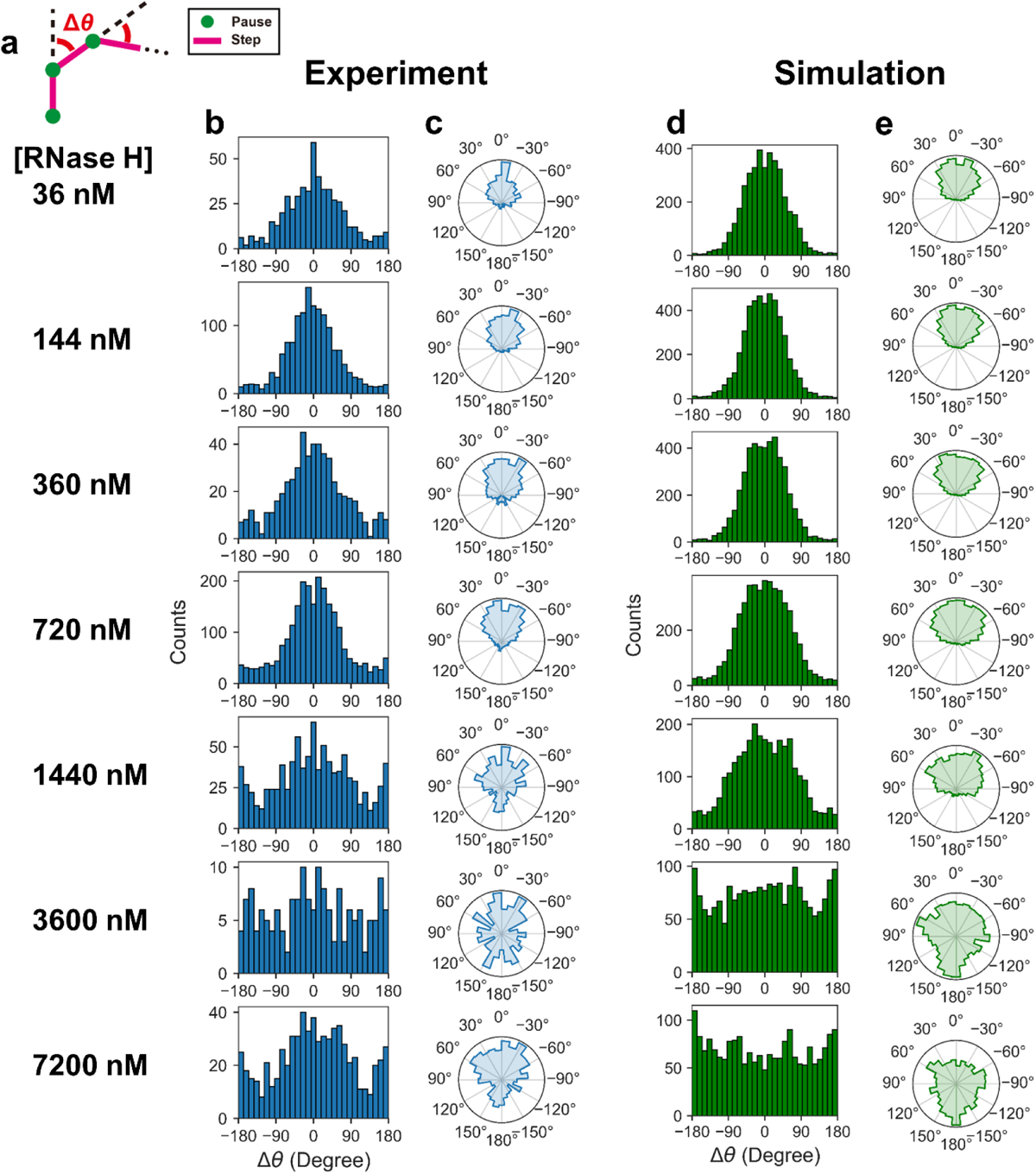
[RNase H] dependence of step angle. **(a)** Schematic illustration describing the calculation of step angle (Δ*θ*). Leftward and rightward steps are defined as positive and negative, respectively. **(b, c)** Histograms (b) and polar plots (c) of step angles at each [RNase H] in the experiments. **(d, e)** Histograms (d) and polar plots (e) of step angles at each [RNase H] in the simulations.

### DNA/RNA sequence with large hybridization rate overcomes trade-off

We summarized relationships between the speed and other performances (Fig. 6, circles). Note that to evaluate the unidirectionality quantitatively, we used the standard deviations of the Δ*θ* distributions (see Methods). [RNase H] dependences of the speed, processivity, run-length, and unidirectionality are also summarized (Supplementary Fig. 19). For both the experiment and simulation, the trade-off between the speed and other performances was clearly observed at high [RNase H].

**Figure 6.**
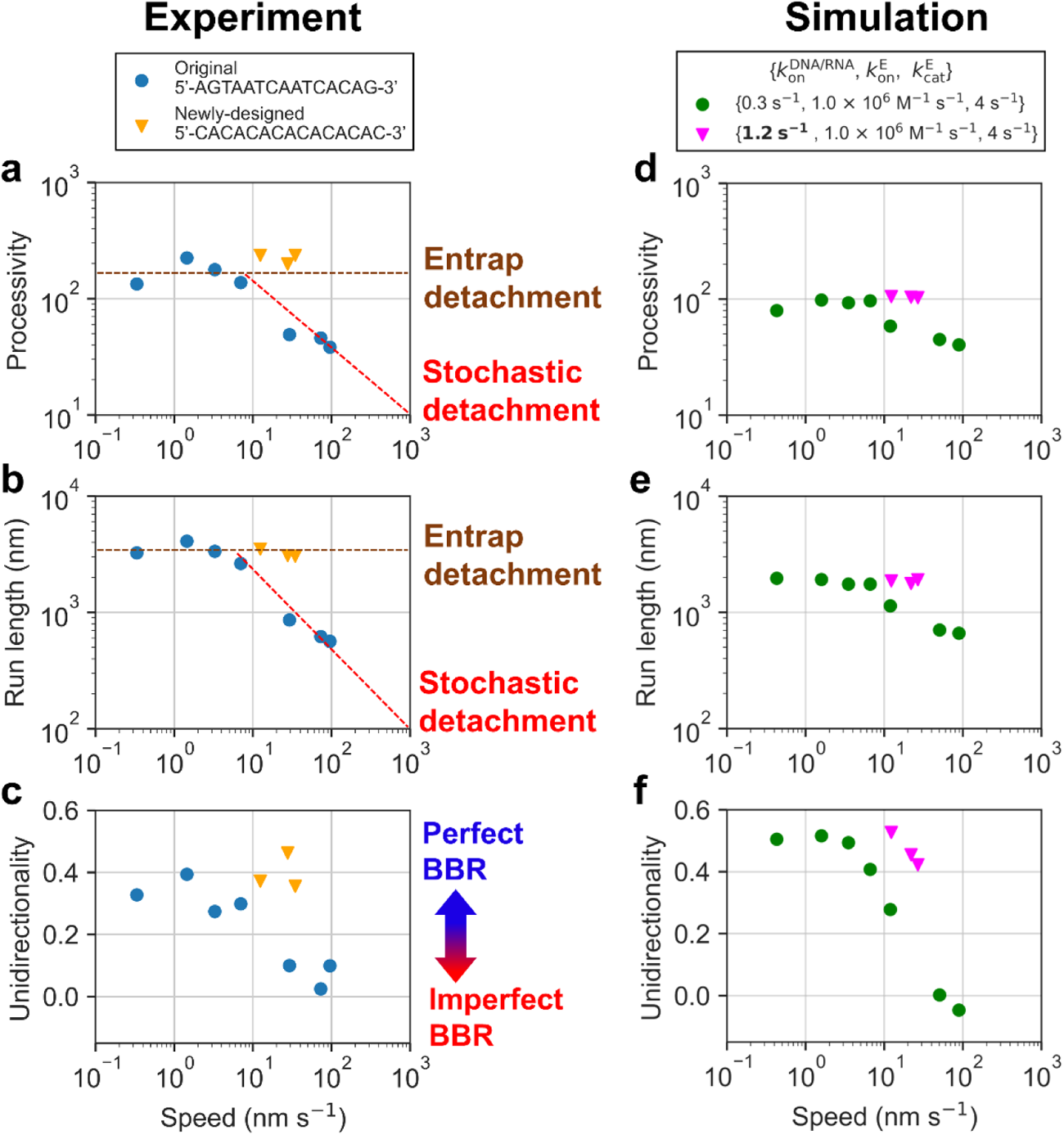
Trade-off between motor speed and other performances. **(a-c)** Relationship between the speed and processivity (a), or run-length (b), or unidirectionality (c) in the experiments. Blue circles indicate the data using original DNA/RNA sequence at 36, 144, 360, 720, 1440, 3600, and 7200 nM RNase H. Orange triangles indicate the data using newly-designed DNA/RNA sequence at 1440, 3600, and 7200 nM RNase H. The upper right and lower left regions of each panel correspond to high and low performances, respectively. Diagonal red dashed lines in (a) and (b) show the trade-off between speed and processivity/run-length due to the stochastic detachment, and horizontal brown dashed lines show the upper limit of processivity/run-length due to the entrap detachment. In (c), high and low unidirectionalities correspond to the perfect and imperfect BBR, respectively. **(d-f)** Relationship between the speed and processivity (d) or run-length (e) or unidirectionality (f) in the simulations. Green circles indicate the data using *k*_on_^DNA/RNA^ = 0.3 s^-1^, k_on_^E^ = 1.0 x 10^6^M^-1^S^-1^ at, 36, 144, 360, 720; 1440, 3600, and 7200 nM RNase H. Pink triangles indicates the data using *k*on^DNA/RNA^ = 1.2 s^-1^, k_on_^E^ = 1.0 x 10^6^ M^-1^S^-1^ and 7200 nM RNase H.

Considering the mechanism shown in Fig. 3, we anticipated that large *k*_on_ ^DNA/RNA^ improves the trade-off by expanding the perfect BBR region (Fig. 3, region (I)) to high [RNase H] region.

Actually, our simulation predicted that 4-times increase in *k*_on_^DNA/RNA^ results in no decreases in processivity, run-length, and unidirectionality even at 7200 nM RNase H, although the speed slightly decreases to 30 nm s^−1^ (Fig. 6d-f, Supplementary Fig. 19e-h, pink triangles, and Supplementary Video 7).

Then, based on the prediction by the simulation, we re-designed the DNA/RNA sequence. We introduced a repetitive sequence and increased GC content, which have been reported to increase *k*_on_ ^DNA/RNA^ (Supplementary Table 1)^30^. Indeed, in the bulk assay, newly-designed sequence showed 3.8-times larger *k*_on_ ^DNA/RNA^ than that of original one, while *k*_off_ ^DNA/RNA^ decreased (Extended Data Fig. 3a, b, e, and f). We also confirmed that *k*_on_^E^ and *k*_cat_^E^ were not largely different from those of the original sequence (Extended Data Fig. 3c, d, g, and h). Then, single-particle tracking of the DNA-AuNP motor with the new sequence was performed at 1440, 3600, and 7200 nM RNase H (Supplementary Fig. 20 and 21). As a result, the trade-off between the speed and other performances was actually improved (Fig. 6a-c, orange triangles). Notably, at 7200 nM RNase H, the newly-designed motor simultaneously achieved speed of 30 nm s^−1^, processivity of 200, run-length of 3 μm, and high unidirectionality (Supplementary Video 8).

## Discussion

Here high-speed/high-precision single-particle tracking resolved the elementary pauses and steps of the DNA-AuNP motor (Fig. 1). Reflecting its nanoscale, the step size (∼20 nm) was comparable to those of motor proteins such as 8 nm step for kinesin-1^12^ and 36 nm step for myosin V^13,31^. Furthermore, the step size was independent on [RNase H], suggesting that it is determined by the particle size. It has been reported that DNA-microparticle motor shows higher speed than that of DNA-nanoparticle motor, despite its small diffusion coefficient^16,17^. Our results suggest that the large particle shows large step size and facilitates the fast motion.

In contrast to the constant step size, the pause length largely shortened as [RNase H] increased (Fig. 2). The geometry-based kinetic simulation quantitatively reproduced the experiment and revealed that RNase H binding to DNA/RNA duplex is the bottleneck at nanomolar RNase H (Fig. 2 and 3). Our result is consistent with previous studies of nano-sized BBR motors where the speed of ∼1 nm s^−1^ were reported at 14–120 nM RNase H^18,29^, and suggests that the speed of these motors can be increased by increasing [RNase H].

At high [RNase H], however, our study also revealed that processivity, run-length, and unidirectionality were sacrificed due to the switch of the bottleneck from RNase H binding to DNA/RNA hybridization (Fig. 4-6). Our simulation predicted that large hybridization rate improves the trade-off, and the newly-designed motor with large *k*_on_ ^DNA/RNA^ simultaneously achieved speed, processivity, and run-length comparable to motor proteins (Fig. 6)^11–14^. Note that the processivity is even higher than that of a biological BBR motor processive chitinase that moves unidirectionally with the speed of 50 nm s^−1^ and processivity of 60∼80 along the single chitin chain on crystalline chitin surface^14,32^.

In our simulation, we can easily change the values of *k*_on_^DNA/RNA^, *k*_on_ ^E^, and *k*_cat_^E^. In general, the larger the values of these kinetic parameters, the better performances are expected. In the experiment, large *k*_on_^DNA/RNA^ can be achieved by re-designing DNA/RNA sequences, as demonstrated in this study (Fig. 6). The large *k*_on_^E^ and *k*_cat_^E^ are more challenging, but possible by using RNase H from other species or by engineering RNase H mutant. Furthermore, our simulation is versatile for other artificial BBR motors such as a protease-modified microparticle motor moving on a peptide-modified substrate recently reported^33^.

Another important issue is the fundamental limitation of processivity and run-length by the entrap detachment (Extended Data Fig. 2), which originates from the fact that the motion of DNA-AuNP motor is not completely unidirectional (Fig. 5). Several studies have reported that microparticle-based BBR motors translocate in a quasi-unidirectional manner along micropatterned fuel surfaces^16,33^. Also, DNA nanotubes and origami tiles, which allow for the elaborate arrangement of interaction sites, can serve as a linear or more complex track for the DNA-AuNP motors^9,34^. Furthermore, nano-sized objects with anisotropic shapes such as DNA origami rod and gold nanorod would restrict the Brownian motion to one direction and improve the unidirectionality^29,35^. Our next goal is development of nano-sized BBR motors capable of directional and speed controls, which are applicable to sensing or computation based on the changes in the motional behaviors^36^.

## Methods

### Regents

DNAs and RNAs used in this study (Supplementary Table 1) were commercially synthesized and purified by high-performance liquid chromatography. DNA and DNA/RNA chimera substrate were purchased from Tsukuba Oligo Service (Ibaraki, Japan). DNA anchor (3’-alkyne-modified) was purchased from Fasmac (Kanagawa, Japan). RNase H from *E. coli* (Product No. 2150A) was purchased from Takara Bio Inc. (Shiga, Japan).

### Preparation of 100 nm DNA-AuNPs

Twenty μL of 100 µM DNA leg T30 in the phosphate-buffered saline (PBS, pH 7.4) was added to 180 μL of a suspension of commercially purchased 100 nm diameter gold colloid (BBI Solutions) in a glass vial. The solution was cooled and frozen by placing it in a –30 °C freezer for 20 min. Then, the solution was allowed to thaw at room temperature for 15 min. Freeze-thaw cycles were repeated three times. Then, the solution was centrifuged (6000 rpm for 1 min) and the pellet was washed with 100 μL of PBS containing Triton X-100 (0.75% v/v) seven times. The obtained DNA-AuNPs were stored at 4 °C.

### Measurement of DNA density on AuNP surface

Ten µL of fluorescein (FAM) labeled DNA-AuNPs (∼10 pM, Supplementary Fig. 1) was mixed with 10 µL of 1M DTT and incubated at 50°C for 30–60 min. After the incubation, the solution was centrifuged (6000 rpm for 1 min) and supernatant was collected. Then, 180 µL of PBS was added and fluorescence intensity at 520 nm (excitation: 490 nm) was measured using a fluorescence spectrometer (FP8300, JASCO). Released FAM-DNA concentration was quantified from the fluorescence intensity using a calibration curve. The number of DNAs on AuNP was calculated from the ratio of the concentration of released FAM-DNA to that of AuNP, and then DNA density on AuNP surface was calculated as the number of DNAs on AuNP divided by the surface area assuming a sphere with a 100 nm diameter.

### Preparation of KOH washed cover glass used for flow cell

The cover glass (9×30 mm, #1, Matsunami Glass Industry, Japan) was washed and sonicated in ethanol for 10 min. Then, the glass was rinsed 3 times with ultrapure water (18.2 MΩ cm^−1^) and sonicated in ultrapure water for 10 min. The cover glass was rinsed again with ultrapure water. After thoroughly drying with N_2_ blow, the cover glass was immersed in the KOH 10 M for 1 hour. Then, the cover glass was well rinsed with ultrapure water. The cleaned cover glass was dried with N_2_ blow and kept on the clean rack in the closed container prior to use.

### Preparation of amino-silane coated cover glass used for flow cell

The cover glass (24×32 mm, #1 SHT, Matsunami Glass Industry) was washed and sonicated in ethanol for 10 min. Then, the glass was rinsed with ultrapure water 6 times and sonicated in ultrapure water for 10 min. The cover glass was then rinsed again with ultrapure water 6 times and immersed in the piranha solution for 1 hour. The cover glass was rinsed and washed well with ultrapure water. Then, the cover glass was rinsed with ethanol and immersed in [3-aminopropyl]-triethoxysilane in ethanol (2% v/v) for 60 min at room temperature. After that, the cover glass was rinsed well with ethanol and cured in an oven at 110 °C for 10 min. The cover glass was kept on the clean rack in the closed container prior to use.

### Flow cell preparation for single-particle tracking experiment

The flow cell for DNA-AuNP motor observation was prepared by using the amino-silane coated glass (bottom side) and the KOH washed cover glass (top side) and a spacer (a double-sided tape, thickness of 0.62 mm) (Supplementary Fig. 2a). The length and width of each flow cell were 9.0 mm and 2.5 mm, respectively, and a grease (Dow Corning Toray) was used to isolate each flow cell. Each flow cell was then incubated with 10 mg/mL azide-peg_4_-NHS (Tokyo Chemical Industry) dissolved in 100 mM NaHCO_3_ solution (pH 9.0) for 1 hour at room temperature and washed with ∼50 μL ultrapure water (Supplementary Fig. 2b). Then, flow cells were washed with 1 M potassium phosphate (pH 7.4), and 30 μL of 20 µM alkyne-modified DNA anchor, 100 µM Tris(3-hydroxypropyltriazolylmethyl)amine (THPTA), 20 µM CuSO_4_ and 2 mM sodium ascorbate, 1 M potassium phosphate (pH 7.4) was added to the flow cell and incubated for 1 hour at room temperature. After that, the flow cell was washed with PBS, and then 30 μL of 1 μM Cy3-RNA/DNA chimera in PBS was added and incubated overnight at room temperature in a box with wet filter paper attached to the inside wall. After incubation, unbound RNAs were washed out with ∼50 μL PBS and stored in the box prior to use.

### Measurement of RNA density on cover glass surface

The flow cell of which the bottom surface was modified with Cy3-RNA/DNA chimera was incubated with 20 µL of 100 µg mL^−1^ RNase A in PBS for 5 min at room temperature (Supplementary Fig. 2c). Then, 20 µL of the reacted solution was collected and added into a cuvette for fluorescence measurement. This procedure was repeated at least three times, to assure that all of the Cy3-RNA/DNA chimera on cover glass surface were cleaved by RNase A. The collected solution was then diluted to a final volume of 300 µL with PBS. The concentration of Cy3-RNA was quantified from the fluorescence intensity at 560 nm (excitation: 510 nm) using a calibration curve obtained with a fluorescence spectrometer (FP8300, JASCO). The total number of RNA molecules was calculated as a product of concentration and volume (300 µL). RNA surface density was calculated from the total number of RNA molecules divided by the area of the bottom surface of the flow cell (9.0 mm × 2.5 mm).

### Single-particle tracking of DNA-AuNP motor with scattering imaging

A total internal reflection dark-field microscope was built as previously described^26,37^. We used the point illumination for observation^26,37^. Observation was conducted at 25 ± 1°C, and scattering signals from the DNA-AuNPs were imaged with a high-speed CMOS camera (Fastcam AX100; Photron). An observation buffer (50 mM Tris-HCl, 10 μM DTT, 0.75% v/v Triton X-100, 3 mM MgCl_2_, pH 8.3) was used. The flow cell was placed on the stage of microscope and incubated with 3 pM DNA-AuNPs solution for 15 min. Then, the flow cell was washed with the observation buffer, and the sample stage was left untouched at least 10 min to minimize the stage drift. The motion of DNA-AuNP was observed with designated RNase H concentration ([RNase H]). Recording rate of 20 frames per second (fps) was used for 36, 144, and 360 nM RNase H, 250 fps for 720 and 1440 nM RNase H, and 1000 fps for 3600 and 7200 nM RNase H. For each recording rate, we adjusted the laser power density to avoid the saturation of the scattering signal (Supplementary Fig. 3). At 36 nM RNase H condition, cooling fan of the CMOS camera had to be turned on for the long-time recording for more than 300 s to avoid overheating. Then, sequential images of individual DNA-AuNP motors were analyzed separately by home-built Python code. At each frame, X- and Y-coordinates of the centroid of each DNA-AuNP motor were determined by the two-dimensional Gaussian fitting. The localization precision at 36 nM RNase H condition was 3.1 nm for both X- and Y-coordinates, and those at other [RNase H] conditions were 0.8–1.0 nm (Supplementary Fig. 3).

### Analysis of step size and pause length

Trajectories of the centroid of the DNA-AuNP motors were analyzed by home-built Python code. First, steps along X-and Y-coordinates were identified separately by using an algorithm developed previously^27^. Then, diagonal steps were counted when the steps along X- and Y-coordinates occurred within ± 5 frames because of the limited accuracy of the step-finding algorithm. Then, steps along X- and Y-coordinates and diagonal steps were combined to obtain all pause lengths and step sizes in entire trajectories.

### Analysis of processivity and run-length

In the single-particle tracking experiments, especially at low [RNase H], significant fraction of the motors did not show detachment from the RNA substrate during observation. Therefore, we followed an approach used for other nano-sized BBR motors^18,29^, in which time course of the fraction of motors remained attached on RNA substrate was measured. Then, time courses were fitted with single exponential decay functions to estimate time constants before detachment (τ_detach_), and processivity and run-length were estimated by following equations, respectively.

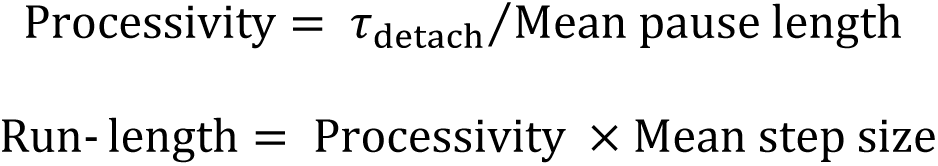

On the other hand, simulation was conducted until each motor showed the detachment. Therefore, for simulation data, we directly estimated the processivity (number of steps before detachment) from each trajectory, and calculated run-length by using the equation described above. For simulation, mean values are calculated from 50 trajectories for each [RNase H] condition.

### Analysis of step angle and unidirectionality

Step angles (Δ*θ*) were calculated from the XY coordinates of three consecutive pauses. In the case of 2D Brownian motion, the histogram of the Δ*θ* converges to a uniform distribution between −180° and 180°, while in the case of completely unidirectional motion, the histogram of the Δ*θ* has a single peak at 0°. Unidirectionality of the DNA-AuNP motor was quantified by using the standard deviation of the Δ*θ* distribution:

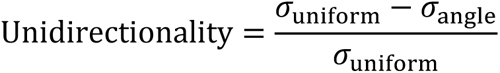

where *σ*_angle_ and *σ*_uniform_ are the standard deviations of the Δ*θ* distribution of the data and that assuming the uniform distribution, respectively. Theoretically, the value of the *σ*_uniform_ is 103.9°, and the upper and lower limits of the unidirectionality are 1 (when *σ*_angle_ is 0°) and 0 (when *σ*_angle_ = *σ*_uniform_), respectively.

### DNA/RNA hybridization and RNA hydrolysis assays in bulk solution

We applied a fluorescence-based assay in which the quantum yield of Cy3-oligonucleotide is sensitive to the hybridization state^38,39^. A fluorescence spectrometer (FP8300, JASCO) was used to monitor hybridization between Cy3-labeled RNA and the complementary ssDNA in a free-solution. All measurement was performed at room temperature. The reaction buffer was the same as that used for the DNA-AuNP motor observation. Fluorescence from 998 μL of 10 nM Cy3-RNA in a cuvette was monitored continuously with excitation and emission wavelengths of 515 and 565 nm and slit widths of 10 and 20 nm, respectively. Two μL of concentrated DNA was added to the 998 μL Cy3-RNA solution while stirring the solution at 500 rpm with a magnetic stirrer. The final concentrations of DNA ([DNA]_0_) were 6–100 nM. Then, the change in fluorescence intensity was monitored for 300 s with 1 s time resolution, and apparent hybridization rate at each [DNA]_0_ was estimated as the inverse of the relaxation time. Measurement was repeated at least 3 times for each condition. The values of *k*_on_ ^DNA/RNA^ and *k*_off_ ^DNA/RNA^ (Supplementary Table 2) were determined from the [DNA]_0_ dependence of the apparent hybridization rate (Extended Data Fig. 1e).

For RNA hydrolysis assay, the condition of fluorescence measurement was the same with hybridization assay described above, except for the time resolution of 0.02 s. Ten nM Cy3 RNA and 20 nM DNA were mixed and incubated for more than 10 min at room temperature for hybridization. Then, 4 uL of concentrated RNase H was added to the cuvette, and final concentrations of RNase H ([RNase H]_0_) were 69-6912 nM. The decay of fluorescence intensity was monitored for 130 s. Measurement was repeated at least 3 times for each condition. The decay curve was fitted by the single exponential decay to determine the apparent RNA hydrolysis rate. The values of *k*_on_ ^E^ and *k*_cat_^E^ (Supplementary Table 2) were determined from the [RNase H]_0_ dependence of the apparent hybridization rate (Extended Data Fig. 1f).

### Geometry-based kinetic simulation of DNA-AuNP motor

The home-built code for the geometry-based kinetic simulation of DNA-AuNP motor was implemented in Python 3.9. Details of the algorithm and the simulation parameters are described in Supplementary Information (Supplementary Note, Supplementary Fig. 6-8, Supplementary Table 3). First, a two-dimensional pixel matrix was defined to model the RNAs on the glass surface. Pixel size was normalized by the DNA density on the AuNP (0.10 molecules nm^−2^). RNAs were randomly distributed with a ratio of the RNA density (0.022 molecules nm^−2^) to the DNA density. Reaction at a single RNA site is assumed to contain three sequential elementary steps: DNA/RNA hybridization, RNase H binding to DNA/RNA duplex, and RNA hydrolysis. The reaction proceeds within the accessible area of DNA with the radius of 28.0 nm (Supplementary Fig. 7). Rate constants of the three elementary steps, *k*_on_^DNA/RNA^, *k*_on_ ^E^, and *k*_cat_^E^ are the simulation parameters. The optimal set of the rate constants was obtained through the simulation-based fitting (see below) and used as the default values. Dwell time for each elementary step at each RNA site is determined by a random sampling from the exponential distribution: P(τ) = *k*exp(−*k*t), and the site with the shortest dwell time is changed to the next state. Simulation steps proceeded in units of reaction events. Constraint of the motor position was considered by introducing mobile region of each DNA/RNA hybrid (25.2 nm) (Supplementary Fig. 8). The motor position was determined randomly within the region where all mobile regions overlap. Unless all mobile regions overlapped, the motor position was fixed. To reproduce the experimental trajectory accurately, Gaussian noise with standard deviation of 2 or 6 nm was added to the raw simulated trajectories (Supplementary Fig. 9 and 10).

### Simulation-based fitting to optimize kinetic parameters

Simulation-based fitting was employed to obtain the optimal set of the rate constants (*k*_on_^DNA/RNA^, *k* ^E^, *k*_cat_^E^) which reproduce the experimental results more precisely (Supplementary Fig. 11-16), because the rate constants on the surface can change from those estimated in the bulk assay (Extended Data Fig. 1, Supplementary Table 2). The geometry-based kinetic simulation of the DNA-AuNP motor was performed with all combinations of *k*_on_^DNA/RNA^ = {0.01, 0.05, 0.1, 0.2, 0.3, 0.4, 0.5, 1.0, 5.0} s^−1^, *k*_on_^E^ = {0.5, 1.0, 1.5, 2.0, 2.5} × 10^6^ M^−1^s^−1^, and *k*_cat_^E^ = {0.5, 1.0, 2.0, 3.0, 4.0, 5.0, 6.0} s^−1^. The [RNase H] of 36, 144, 360, 720, 1440, 3600, and 7200 nM were tried for each parameter set, and 20 simulations were carried out for each combination. Pause length and step size were calculated from the simulated trajectories by a step-finding algorithm. As the evaluation function, the sum of the mean squared errors (MSE) of the pause length and step size against the experimental data was calculated by using following equation:

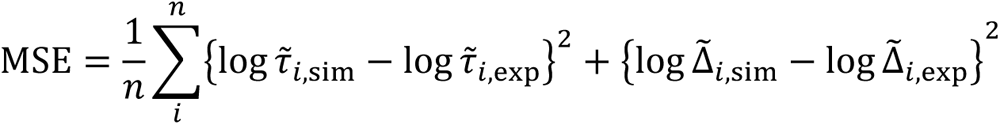

where 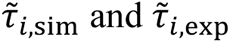 are the median pause length in simulation and experiment, Δ^-^_i,sim_ and Δ^-^_i,exp_ are the median step sizes in simulation and experiment, and *n* is the number of the conditions of [RNase H]. The parameter set (*k*_on_ ^DNA/RNA^, *k*_on_^E^, *k*_cat_^E^) which showed the minimal MSE was used for the subsequent 50 simulations (Supplementary Figure 16, Supplementary Table 3).

---

**Extended Data Fig. 1.**
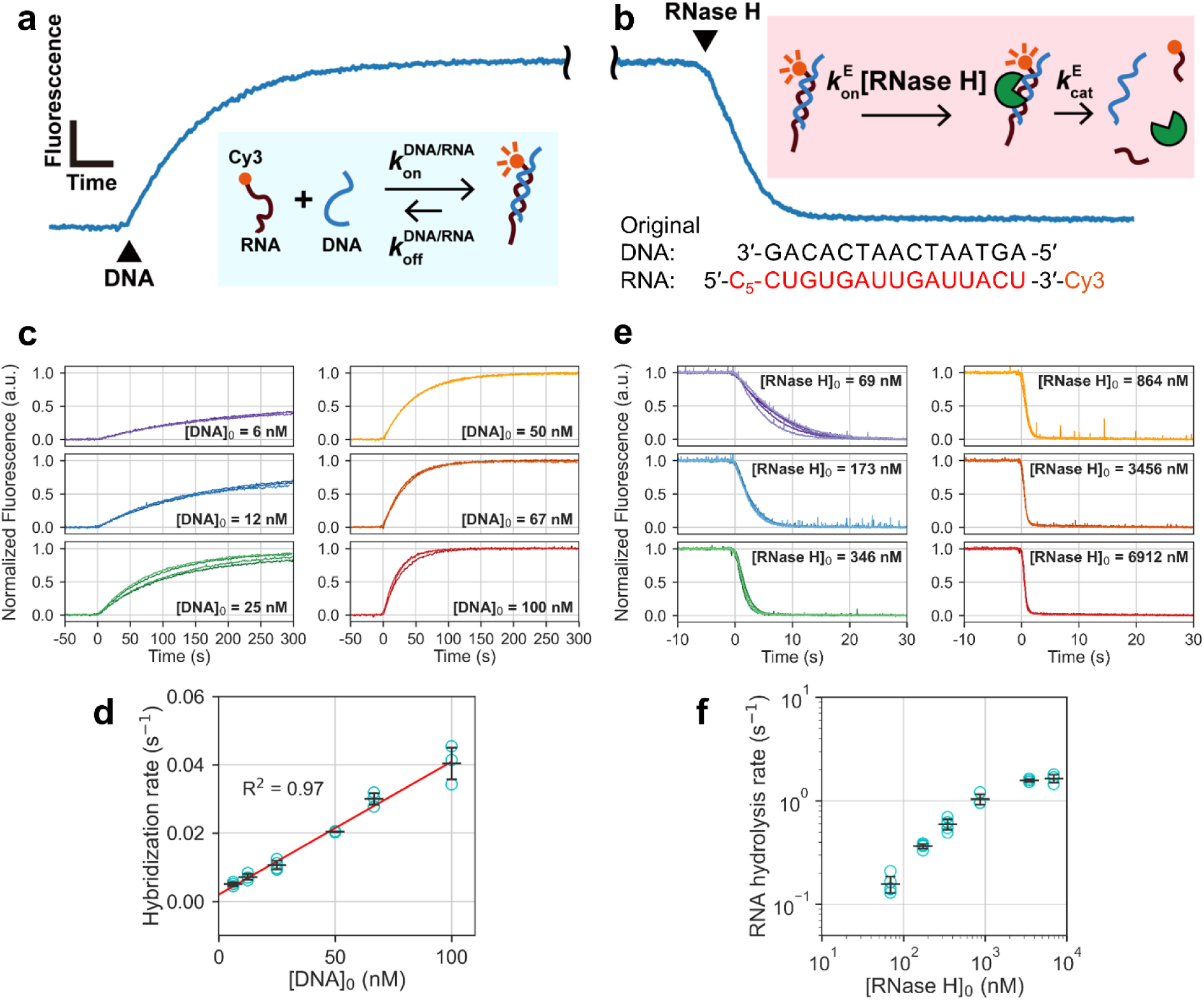
Determination of kinetic rate constants in bulk solution. **(a, b)** Schematic illustration and kinetic model of (a) DNA/RNA hybridization and (b) RNA hydrolysis assays. Cy3-labeled RNA was used as a probe of DNA/RNA hybridization. The same sequence to that of the DNA/RNA duplex part of the original sequence and the same solution composition for single-particle tracking experiments were used. **(c)** Normalized time courses of fluorescence intensity of the DNA/RNA hybridization assay. Initial RNA concentration was 10 nM, and measurements were carried out at least in triplicate for each initial DNA concentration ([DNA]_0_). Error bars represent s.d. Time resolution was 1 s, and excitation and emission wavelengths were set at 510 nm and 560 nm, respectively. **(d)** Plots of apparent hybridization rate as a function of [DNA]_0_. The values of *k_on_ ^DNA/RNA^* and *k_off_^DNA/RNA^* were determined as the slope and intercept of the linear fitting (red line), respectively. **(e)** Normalized time courses of fluorescence intensity of the RNA hydrolysis assay. Initial DNA and RNA concentrations were 10 nM, and measurements were carried out at least in triplicate for each RNase H concentration ([RNase H]_0_). Time resolution was 0.02 s, and same excitation and emission wavelengths were used as (c). **(f)** Plots of apparent hydrolysis rates as a function of [RNase H]_0_. Error bars represent s.d. The value of *k*_on_^E^ was calculated by dividing the rates with [RNase H]_0_ at 69 and 173 nM RNase H. The value of *k*_cat_^E^ was estimated from the saturating hydrolysis rate at 3456 and 6912 nM RNase H.

**Extended Data Fig. 2.**
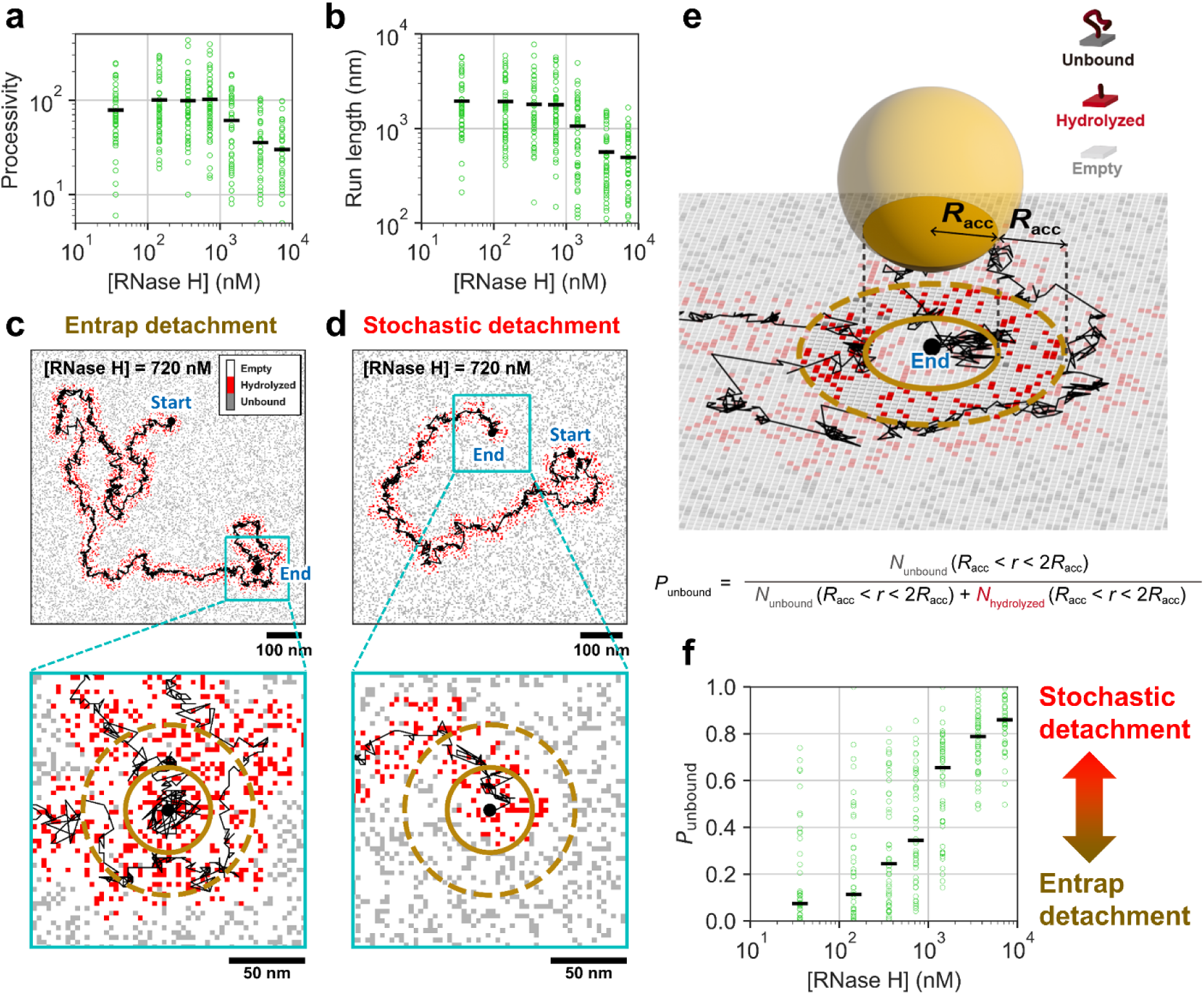
[RNase H] dependence of processivity and run-length, and two different mechanisms of detachment revealed by the simulation. **(a, b)** Plots for processivity (a) and run-length (b) at different [RNase H] conditions in the simulations. Mean values are calculated from 50 trajectories for each condition (black bars). **(c, d)** Two representative trajectories which show entrap (c) and stochastic (d) detachment at 720 nM RNase H. Black lines correspond to the trajectories of the centroid of the motors. Large black dots indicate the start and end of each trajectory. Regions around the end points are indicated by cyan squares and enlarged in lower panels. Dark yellow solid and dashed circles indicate the accessible region at the final frame and the region with the twice radius of the accessible region (2×*R*_acc_), respectively. Trajectories are overlaid on the distribution of RNA states on the surface. White, red, and gray dots correspond to the empty, hydrolyzed, and unbound RNA sites, respectively. **(e)** An example of 3D view corresponding to the last frame before entrap detachment. The *P*_unbound_ is defined as the ratio of unbound RNA sites to unbound and hydrolyzed RNA sites in the doughnut region with the radius between *R*_acc_ to 2×*R*_acc_. Small or large value of the *P*_unbound_ corresponds to entrap or stochastic detachment is major, respectively. **(f)** [RNase H] dependence of the *P*_unbound_. Median values are calculated for each condition and shown by black horizontal bars.

**Extended Data Fig. 3.**
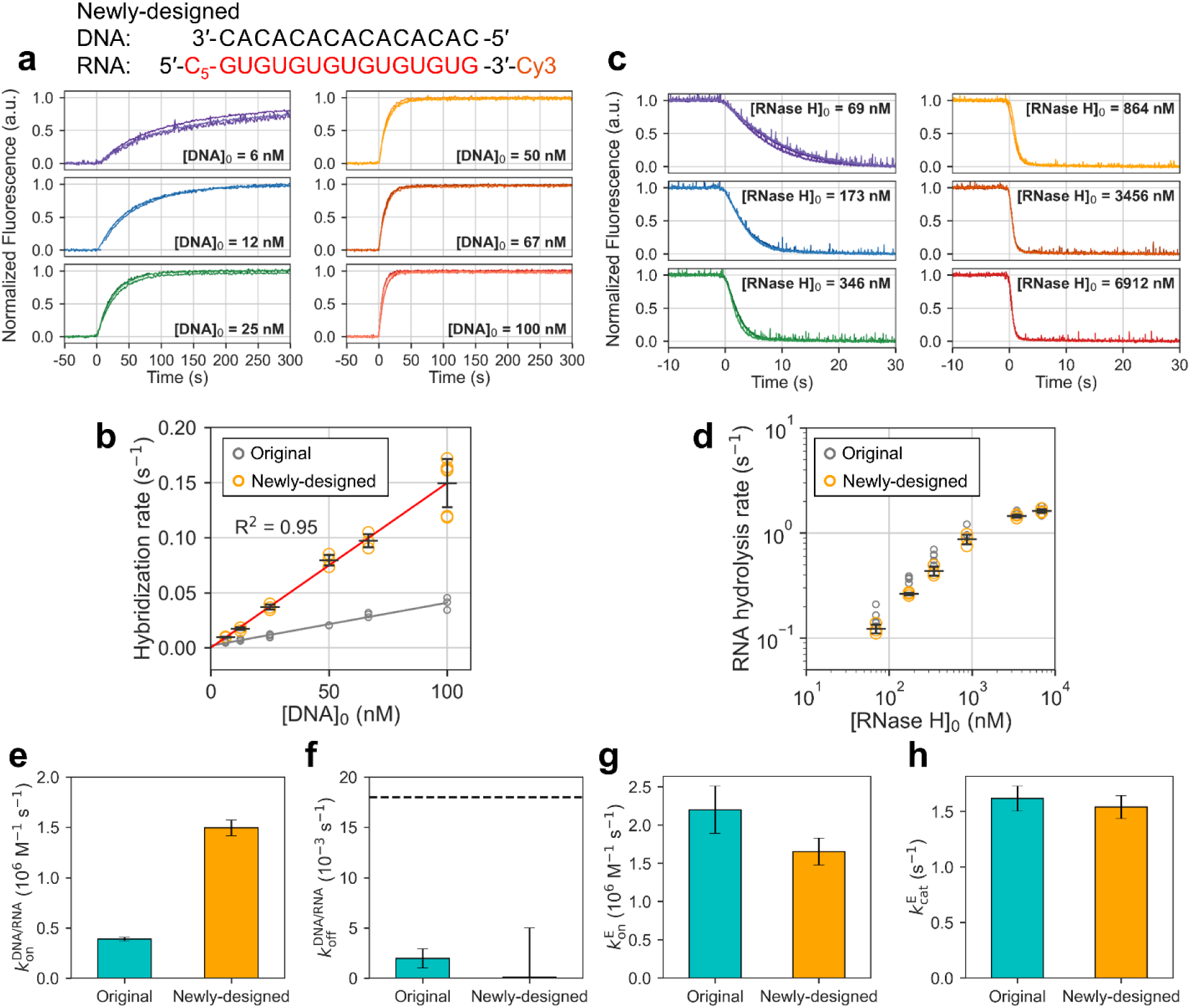
Determination of kinetic rate constants in bulk solution with newly-designed DNA/RNA sequence. **(a)** Normalized time courses of fluorescence intensity of the DNA/RNA hybridization assay. Initial RNA concentration was 10 nM, and measurements were carried out at least in triplicate for each initial DNA concentration ([DNA]_0_). ― Error bars represent s.d. Time resolution was 1 s, and excitation and emission wavelengths were set at 510 nm and 560 nm, respectively. **(b)** Plots of apparent hybridization rate as a function of [DNA]_0_. The values of *k*_on_^DNA/RNA^ and *k*_off_^DNA/RNA^ were determined as the slope and intercept of the linear fitting (red line), respectively. **(c)** Normalized time courses of fluorescence intensity of the RNA hydrolysis assay. Initial DNA and RNA concentrations were 10 nM, and measurements were carried out at least in triplicate for each RNase H concentration ([RNase H]_0_). Time resolution was 0.02 s, and same excitation and emission wavelengths were used as (a). **(d)** Plots of hydrolysis rates as a function of [RNase H]_0_. Error bars represent s.d. **(e-h)** Bar graphs of rate constants for comparison between original (cyan) and designed (orange) DNA/RNA sequences. Error bars represent s.d. Black dashed line in (f) indicates the minimum pause-step cycle rate in single-particle tracking experiments calculated by 1/median pause length at 36 nM RNase H.

## ASSOCIATED CONTENT

### Supporting Information

The Supporting Information is available free of charge.

## AUTHOR INFORMATION

### Author Contributions

R.I. conceived the research. T.H., A.O., and R.I. designed the experiments. T.H. performed sample preparations, biochemical and fluorescence imaging-based characterizations, and single-particle tracking. T.H. and A.O. performed the data analysis. T.H. developed geometry-based kinetic simulation. R.I. supervised the project and preparing the manuscript with T.H. All authors discussed the results.

### Funding Sources

MEXT | Japan Society for the Promotion of Science (JSPS) - 23K13645, 24H01732 [Harashima] Tsugawa Foundation. [Harashima]

MEXT | Japan Society for the Promotion of Science (JSPS) - 18H05424, 23H04434 [Iino]

### Notes

The authors declare no competing financial interest.

## ACKNOWLEDGMENT

The authors thank Yasuko Okuni, Yayoi Kon, and Mayuko Yamamoto for their technical assistance in sample preparations, Kosuke Matsumoto and Rosie Graham for helpful discussions.

## REFERENCES

1. Erbas-Cakmak, S., Leigh, D. A., McTernan, C. T. & Nussbaumer, A. L. Artificial Molecular Machines. Chem. Rev. 115, 10081–10206, (2015).

2. Kassem, S. et al. Artificial molecular motors. Chem. Soc. Rev. 46, 2592–2621, (2017).

3. Li, S. et al. A DNA nanorobot functions as a cancer therapeutic in response to a molecular trigger in vivo. Nat. Biotechnol. 36, 258–264, (2018).

4. Chen, Y. J., Groves, B., Muscat, R. A. & Seelig, G. DNA nanotechnology from the test tube to the cell. Nat. Nanotechnol. 10, 748–760, (2015).

5. Iino, R., Kinbara, K. & Bryant, Z. Introduction: Molecular Motors. Chem. Rev. 120, 1–4, (2020).

6. Lund, K. et al. Molecular robots guided by prescriptive landscapes. Nature 465, 206–210, (2010).

7. Gu, H., Chao, J., Xiao, S. J. & Seeman, N. C. A proximity-based programmable DNA nanoscale assembly line. Nature 465, 202–205, (2010).

8. Yin, P., Yan, H., Daniell, X. G., Turberfield, A. J. & Reif, J. H. A unidirectional DNA walker that moves autonomously along a track. Angew. Chem. Int. Ed. 43, 4906–4911, (2004).

9. Wickham, S. F. et al. A DNA-based molecular motor that can navigate a network of tracks. Nat. Nanotechnol. 7, 169–173, (2012).

10. Pan, J. et al. Visible/near-infrared subdiffraction imaging reveals the stochastic nature of DNA walkers. Sci. Adv. 3, e1601600, (2017).

11. Imai, H. et al. Direct observation shows superposition and large scale flexibility within cytoplasmic dynein motors moving along microtubules. Nat. Commun. 6, 8179, (2015).

12. Schnitzer, M. J. & Block, S. M. Kinesin hydrolyses one ATP per 8-nm step. Nature 388, 386–390, (1997).

13. Ali, M. Y. et al. Myosin V is a left-handed spiral motor on the right-handed actin helix. Nat. Struct. Mol. Biol. 9, 464–467, (2002).

14. Nakamura, A., Okazaki, K. I., Furuta, T., Sakurai, M. & Iino, R. Processive chitinase is Brownian monorail operated by fast catalysis after peeling rail from crystalline chitin. Nat. Commun. 9, 3814, (2018).

15. Cha, T. G. et al. A synthetic DNA motor that transports nanoparticles along carbon nanotubes. Nat. Nanotechnol. 9, 39–43, (2014).

16. Yehl, K. et al. High-speed DNA-based rolling motors powered by RNase H. Nat. Nanotechnol. 11, 184–190, (2016).

17. Blanchard, A. T. et al. Highly Polyvalent DNA Motors Generate 100+ pN of Force via Autochemophoresis. Nano Lett. 19, 6977–6986, (2019).

18. Bazrafshan, A. et al. DNA Gold Nanoparticle Motors Demonstrate Processive Motion with Bursts of Speed Up to 50 nm Per Second. ACS Nano 15, 8427–8438, (2021).

19. Ron, M. & Rob, P. Cell Biology by the Numbers (Garland Science, 2015).

20. Howard, J. Mechanics of motor proteins and the cytoskeleton (Sinauer Associates, 2001).

21. Isojima, H., Iino, R., Niitani, Y., Noji, H. & Tomishige, M. Direct observation of intermediate states during the stepping motion of kinesin-1. Nat. Chem. Biol. 12, 290–297, (2016).

22. Nishiyama, M., Higuchi, H. & Yanagida, T. Chemomechanical coupling of the forward and backward steps of single kinesin molecules. Nat. Cell. Biol. 4, 790–797, (2002).

23. Nishizaka, T. et al. Chemomechanical coupling in F1-ATPase revealed by simultaneous observation of nucleotide kinetics and rotation. Nat. Struct. Mol. Biol. 11, 142–148, (2004).

24. Wolff, J. O. et al. MINFLUX dissects the unimpeded walking of kinesin-1. Science 379, 1004–1010, (2023).

25. Ueno, H. et al. Simple dark-field microscopy with nanometer spatial precision and microsecond temporal resolution. Biophys. J. 98, 2014–2023, (2010).

26. Ando, J. et al. Single-Nanoparticle Tracking with Angstrom Localization Precision and Microsecond Time Resolution. Biophys. J. 115, 2413–2427, (2018).

27. Kerssemakers, J. W. et al. Assembly dynamics of microtubules at molecular resolution. Nature 442, 709–712, (2006).

28. Zhang, L., Piranej, S., Namazi, A., Narum, S. & Salaita, K. “Turbo-Charged” DNA Motors with Optimized Sequence Enable Single-Molecule Nucleic Acid Sensing. Angew. Chem. Int. Ed. 63, e202316851, (2024).

29. Bazrafshan, A., et al. Tunable DNA Origami Motors Translocate Ballistically Over μm Distances at nm/s Speeds. Angew. Chem. Int. Ed. 59, 9514–9521, (2020).

30. Hertel, S. et al. The stability and number of nucleating interactions determine DNA hybridization rates in the absence of secondary structure. Nucleic Acids Res. 50, 7829–7841, (2022).

31. Mehta, A. D. et al. Myosin-V is a processive actin-based motor. Nature 400, 590–593, (1999).

32. Nakamura, A. et al. Rate constants, processivity, and productive binding ratio of chitinase A revealed by single-molecule analysis. Phys. Chem. Chem. Phys. 20, 3010–3018, (2018).

33. Korosec, C. S. et al. Motility of an autonomous protein-based artificial motor that operates via a burnt-bridge principle. Nat. Commun. 15, 1511, (2024).

34. Ibusuki, R. et al. Programmable molecular transport achieved by engineering protein motors to move on DNA nanotubes. Science 375, 1159–1164, (2022).

35. Zhou, C., Duan, X. Y. & Liu, N. A plasmonic nanorod that walks on DNA origami. Nat. Commun. 6, (2015).

36. Piranej, S., Bazrafshan, A. & Salaita, K. Chemical-to-mechanical molecular computation using DNA-based motors with onboard logic. Nat. Nanotechnol. 17, 514–523, (2022).

37. Ando, J. et al. Small stepping motion of processive dynein revealed by load-free high-speed single-particle tracking. Sci. Rep. 10, 1080, (2020).

38. Sanborn, M. E., Connolly, B. K., Gurunathan, K. & Levitus, M. Fluorescence properties and photophysics of the sulfoindocyanine Cy3 linked covalently to DNA. J. Phys. Chem. B 111, 11064–11074, (2007).

39. Peterson, E. M., Manhart, M. W. & Harris, J. M. Single-Molecule Fluorescence Imaging of Interfacial DNA Hybridization Kinetics at Selective Capture Surfaces. Anal. Chem. 88, 1345–1354, (2016).

