## Supplementary Information for "Mechanism-based design of DNA-nanoparticle motor with high speed and processivity comparable to motor proteins"

**Contents**

**Supplementary Note S4**

**Supplementary Table 1. Nucleotide sequences of DNA and RNA used in this study. S7**

**Supplementary Table 2. Kinetic parameters estimated by the fluorescence bulk assays S8**

**Supplementary Table 3. Geometric and kinetic parameters used in the simulation S8**

**Supplementary Figure 1. Surface modification of AuNP with DNA and characterization S9**

**Supplementary Figure 2. Modification of glass substrate with RNA and characterization S10**

**Supplementary Figure 3. Data recording conditions and localization precisions S11**

**Supplementary Figure 4. Typical trajectories of experiment and simulation at each [RNase H] S12**

**Supplementary Figure 5. Distributions of pause length, step size, and speed at each [RNase H] for the experimental data S13**

**Supplementary Figure 6. Flowchart of geometry-based kinetic simulation S14**

**Supplementary Figure 7. Geometric model to calculate the accessible radius S15**

**Supplementary Figure 8. Configuration of the mobile region for the motion of DNA-AuNP motor S16**

**Supplementary Figure 9. Optimization of noise added to simulation trajectory to reproduce experimental trajectory with camera fan on S17**

**Supplementary Figure 10. Optimization of noise added to simulation trajectory to reproduce experimental trajectory with camera fan off S18**

**Supplementary Figure 11. [RNase H] dependence of pause length (a) and step size (b) in kinetic simulations with $k_{on}^{E}$ = 2.5 × $10^6$ $M^{-1}s^{-1}$ S19**

**Supplementary Figure 12. [RNase H] dependence of pause length (a) and step size (b) in kinetic simulations with $k_{on}^{E}$ = 2.0 × $10^6$ $M^{-1}s^{-1}$ S20**

**Supplementary Figure 13. [RNase H] dependence of pause length (a) and step size (b) in kinetic simulations with $k_{on}^{E}$ = 1.5 × $10^6$ $M^{-1}s^{-1}$ S21**

**Supplementary Figure 14. [RNase H] dependence of pause length (a) and step size (b) in kinetic simulations with $k_{on}^{E}$ = 1.0 × $10^6$ $M^{-1}s^{-1}$ S22**

**Supplementary Figure 15. [RNase H] dependence of pause length (a) and step size (b) in kinetic simulations with $k_{on}^{E}$ = 0.5 × $10^{6}$ $M^{-1}s^{-1}$** **S23**

**Supplementary Figure 16. Optimization of rate constants by simulation-based fitting** **S24**

**Supplementary Figure 17. MSD analysis** **S25**

**Supplementary Figure 18. [RNase H] dependence of time-course of the number of RNA sites, X- and Y-coordinates, and step angles (Δ*θ*) in the simulation** **S26**

**Supplementary Figure 19. Summary of [RNase H] dependence of motor performance** **S27**

**Supplementary Figure 20. Step analysis of the DNA-AuNP motor with large hybridization rates** **S28**

**Supplementary Figure 21. Step angle analysis of the DNA-AuNP motor with large hybridization rates** **S29**

**Supplementary References** **S30**

**Supplementary Note**

***Geometric models to calculate accessible radius and mobile region used for the simulation***

A geometric model was constructed to determine the accessible radius ($R_{acc}$) used in the simulation (Supplementary Fig. 7). This model assumes that each single-stranded DNA (ssDNA) on AuNP surface can access to RNA substrate surface within a range shorter than the end-to-end distance of ssDNA ($L_0^{ssDNA}$). Based on the worm-like chain model, the mean-squared end-to-end length of ssDNA can be calculated from the following equation:

$$L_0^{\mathrm{ssDNA}^2} = 2L_\mathrm{p}^{\mathrm{ssDNA}^2}\left[\exp\left(-\frac{L_\mathrm{c}^{\mathrm{ssDNA}}}{L_\mathrm{p}^{\mathrm{ssDNA}}}\right) - 1 + \frac{L_\mathrm{c}^{\mathrm{ssDNA}}}{L_\mathrm{p}^{\mathrm{ssDNA}}}\right]$$

where $L_p^{ssDNA}$ and $L_c^{ssDNA}$ are the persistence and contour lengths of ssDNA, respectively. Following the previous studies[1,2], the $L_p^{ssDNA}$ of 2.0 nm and $L_c^{ssDNA}$ of 0.34 nm per base were used. The value of $L_0^{ssDNA}$ consisting of 45 bases is calculated to be 7.3 nm (Supplementary Fig. 7a). Then, $R_{acc}$ was calculated by using the Pythagorean theorem:

$$R_{\mathrm{acc}}{}^2 = \left(R_{\mathrm{particle}} + L_0^{\mathrm{ssDNA}}\right)^2 - R_{\mathrm{particle}}{}^2$$

where $R_{particle}$ is the radius of the AuNP (50 nm). Then, the $R_{acc}$ was calculated to be 28.0 nm (Supplementary Fig. 7b).

The mobile region of the DNA-AuNP motor in the simulation was determined by considering how long the DNA/RNA bound site can stretch (Supplementary Fig. 8). First, we assumed that the translocation of the DNA-AuNP motor relies on the rolling motion in a lateral direction, and the DNA and RNA are stretched when rolling. The mobile region of each bound site was calculated as the area where the it is not stretched beyond the contour length. The bound site is composed of ssDNA, dsDNA, ssRNA, and RNA/DNA duplex (Supplementary Fig. 8a). According to the literatures[2-4], the total end-to-end distance ($L_0^{total}$) and total contour length ($L_c^{total}$) were calculated to be 20.6 nm and 26.9 nm, respectively. The length of the bound site when the motor is located exactly top of that site was assumed to be $L_0$ (Supplementary Fig. 8b). The relationship between the lateral displacement ($\Delta R$) and the extension ($\Delta L$) after the rolling of the motor is shown in Supplementary Fig. 8c and d. The maximum extension ($\Delta L_{max}$) was calculated to be 6.3 nm from the difference between $L_c^{total}$ and $L_0^{total}$. Then, the maximum value of $\Delta R$ ($\Delta R_{max}$) was calculated to be 25.2 nm, corresponding to the radius of the mobile region for each bound site. In the simulations, the XY-coordinates of the DNA-AuNP are randomly varied within the overlapped region among the mobile regions of all RNA sites bound with DNA. If there is no overlap region, the motor keeps same position. Therefore, the motor is immobile when bound sites are formed throughout the accessible area, and mobile when the bound sites are localized in the small region inside the accessible area.

***Optimization of noise added to simulation trajectory to reproduce experiments***

To precisely reproduce the experimental noise in the simulation, distribution of the standard deviations of X- and Y-coordinates during individual pauses of the DNA-AuNP motor were investigated after the step finding analysis of the experimental data (Supplementary Fig. 9a). Since the cooling fan of the CMOS camera was turned on for the observation at 36 nM RNase H, the standard deviations during the pauses were analyzed separately from other [RNase H] conditions (Supplementary Fig. 9b). The median values for X- and Y-coordinates were calculated to be 6.1 and 6.0 nm, respectively. Then, we optimized the noise added to the raw simulation trajectories. We added the Gaussian noises with width ($\sigma_{\text{noise}}$) of 3, 4, 5, 6, or 7 nm to the simulation trajectories, and compared the standard deviations of the X- and Y-coordinates during the pauses with those of the experiments (Supplementary Fig. 9c and d). Consequently, we concluded that $\sigma_{\text{noise}}$ of 6 nm was the best to reproduce the experimental results. A comparison between experimental and simulated trajectories after the addition of the Gaussian noise is shown in Supplementary Fig. 4.

Next, as the typical condition of single-particle tracking experiments with camera fan off, we analyzed trajectories at 720 nM RNase H. The median value for both X- and Y-coordinates was calculated to be 4.0 nm (Supplementary Fig. 10a), which was smaller than that at 36 nM RNase H with camera fan on. Then, we optimized the noise to reproduce the experimental trajectories. We added the Gaussian noises with $\sigma_{\text{noise}}$ of 1, 2, 3, or 4 nm to the simulation trajectories, and analyzed by the step-finding algorithm. Then, the standard deviations of the X- and Y-coordinates during pauses were compared with those of the experiments (Supplementary Fig. 10b). Consequently, we concluded that $\sigma_{\text{noise}}$ of 2 nm deviation was the best to reproduce the experimental results (Supplementary Fig. 10c). Comparisons between experimental and simulated trajectories after the addition of the Gaussian noise is shown in Supplementary Fig. 4.

***Analysis of mean square displacement (MSD)***

To characterize the motional mode of the DNA-AuNP motor, we analyzed the mean square displacement (MSD) of the trajectory (Supplementary Fig. 17). We calculated the MSD averaged for both time and ensemble according to the following equation:

$$\mathrm{MSD}(\Delta t) = \frac{\sum_{i=0}^{N}\sum_{j=0}^{w_i(\Delta t)}\left[\{x_i(t_j+\Delta t)-x_i(t_j)\}^2+\{y_i(t_j+\Delta t)-y_i(t_j)\}^2\right]}{\sum_{i=0}^{N} w_i(\Delta t)}$$

where $i$ is the identifier of the particles. $j$ is the identifier of the time points. $w_i(\Delta t)$ is the number of data to calculate MSD with $\Delta t$ for $i$th particle. $N$ is the total number of particles. Generally, the MSD scales with Δt as MSD ~ $\Delta t^{\alpha}$, where α is the scaling (or anomalous) factor which characterizes the type of the diffusion[5,6]: $0 < \alpha < 1$ for sub-diffusion, $\alpha = 1$ for simple Brownian diffusion, $1 < \alpha < 2$ for super-diffusion, $\alpha = 2$ for ballistic motion.

All the trajectories of experiments and simulations were used to calculate the MSDs for different [RNase H]s (Supplementary Fig. 17a, d). As results, all MSD plots showed gradual increases in the slopes as the $\Delta t$ increased. This means the gradual increase in the α as $\Delta t$ increased, which suggests the changes in the motional mode of the DNA-AuNP motor depending on the length of $\Delta t$. The shift of MSD curves toward the left (small $\Delta t$) as the increase in [RNase H] is attributed to increase in speed described in Fig. 2. Then, to clarify the relationship between the change in motional mode and pause length depending on [RNase H], the MSD curves were normalized by the median pause length at each [RNase H] (Supplementary Fig. 17b, e). We also calculated time-dependent α from the slope of the MSD curve separated by limited time windows (Supplementary Fig. 17c, f). As $\Delta t$ increased, α increased from ~0.4 (sub-diffusion) to ~1.5 (super-diffusion). Notably, the value of α crossed over 1 around the normalized $\Delta t$ close to 1. This result strongly suggests that the pauses and steps observed in this study cause the sub- and super-diffusions, respectively. Small $\Delta t$ corresponds to the time during single pauses where the large numbers of RNA/DNA duplex immobilize the motor and results in sub-diffusion. On the other hand, large Δt corresponds to the long time period including the multiple pauses and steps, where the translocation of the motor is biased to forward direction due to the burnt-bridge mechanism and results in super-diffusion. The super-diffusive behavior in the long-time period is consistent with the previous studies of the DNA micro- and nano-particle motors[7,8]. Furthermore, at high [RNase H], the decrease in the maximum value of the α was observed. This result is consistent with the decrease in the unidirectionality at high [RNase H] (Fig. 5).

**Supplementary Table 1. Nucleotide sequences of DNA and RNA used in this study**

Nucleotide sequences are displayed in a 5’ to 3’ orientation, the red text indicates RNA bases. The 3’ and 5’ DNA and RNA modifications indicated in the table are illustrated below it.

| ID | Sequence(5'-3') |
|---|---|
| DNA anchor<br>(for cover glass modification) | GAGAGAGATGGGTGCTTTTTTTTTTTTTTT/3'-Alkyne/ |
| Cy3-RNA/DNA chimera<br>Original sequence<br>(for cover glass modification) | GCACCCATCTCTCTCrCrCrCrCrC<br>rCrUrGrUrGrArUrUrGrArUrUrArCrU/3'-Cy3/ |
| Cy3-RNA/DNA chimera<br>Newly-designed sequence<br>(for cover glass modification) | GCACCCATCTCTCTCrCrCrCrCrC<br>rUrGrUrGrUrGrUrGrUrGrUrGrUrGrU/3'-Cy3/ |
| DNA leg T30<br>Original sequence<br>(for AuNP modification) | ThiolC3-5'/TTTTTTTTTTTTTTTTTTTTTTTTTTTTTT<br>AGTAATCAATCACAG/3' |
| FAM-DNA leg T30<br>Original sequence<br>(for AuNP modification) | ThiolC3-5'/TTTTTTTTTTTTTTTTTTTTTTTTTTTTTT<br>AGTAATCAATCACAG/3'-6-FAM |
| DNA leg T30<br>Newly-designed sequence<br>(for AuNP modification) | ThiolC3-5'/TTTTTTTTTTTTTTTTTTTTTTTTTTTTTT<br>ACACACACACACACA/3' |

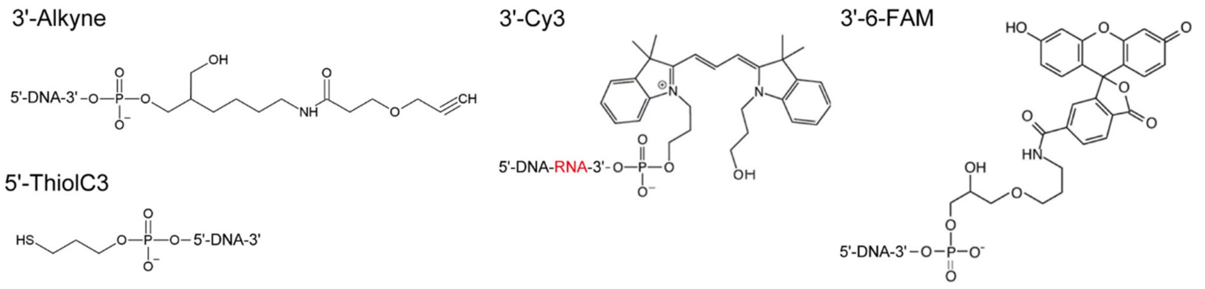

**Supplementary Table 2. Kinetic parameters estimated by the fluorescence bulk assays**

| Sequence | Rate constant | Value |
|---|---|---|
| Original | $k_{on}^{DNA/RNA}$ | $(3.9 \pm 0.5) \times 10^5$ $M^{-1}s^{-1}$ |
| | $k_{off}^{DNA/RNA}$ | $(2.0 \pm 0.9) \times 10^{-3}$ $s^{-1}$ |
| | $k_{on}^{E}$ | $(2.2 \pm 0.3) \times 10^6$ $M^{-1}s^{-1}$ |
| | $k_{cat}^{E}$ | $1.6 \pm 0.1$ $s^{-1}$ |
| Newly-designed | $k_{on}^{DNA/RNA}$ | $(1.5 \pm 0.1) \times 10^6$ $M^{-1}s^{-1}$ |
| | $k_{off}^{DNA/RNA}$ | $(0.1 \pm 5.0) \times 10^{-3}$ $s^{-1}$ |
| | $k_{on}^{E}$ | $(1.7 \pm 0.2) \times 10^6$ $M^{-1}s^{-1}$ |
| | $k_{cat}^{E}$ | $1.5 \pm 0.1$ $s^{-1}$ |

**Supplementary Table 3. Geometric and kinetic parameters used in the simulation**

| **Parameter** | **Default value** |
|---|---|
| Particle diameter | 100 nm |
| DNA surface density | 0.1 molecules $nm^{-2}$ ($1.0 \times 10^5$ molecules $\mu m^{-2}$) |
| RNA surface density | 0.022 molecules $nm^{-2}$ ($2.2 \times 10^4$ molecules $\mu m^{-2}$) |
| Array size (pixel) | 2000×2000 |
| Array size (nm) | 6324 nm × 6324 nm |
| Accessible radius, $R_{acc}$ | 28.0 nm |
| Mobile radius, $R_{mobile}$ | 25.2 nm |
| $k_{on}^{DNA/RNA}$ | 0.3 $s^{-1}$ |
| $k_{on}^{E}$ | $1.0 \times 10^6$ $M^{-1}s^{-1}$ |
| $k_{cat}^{E}$ | 4.0 $s^{-1}$ |

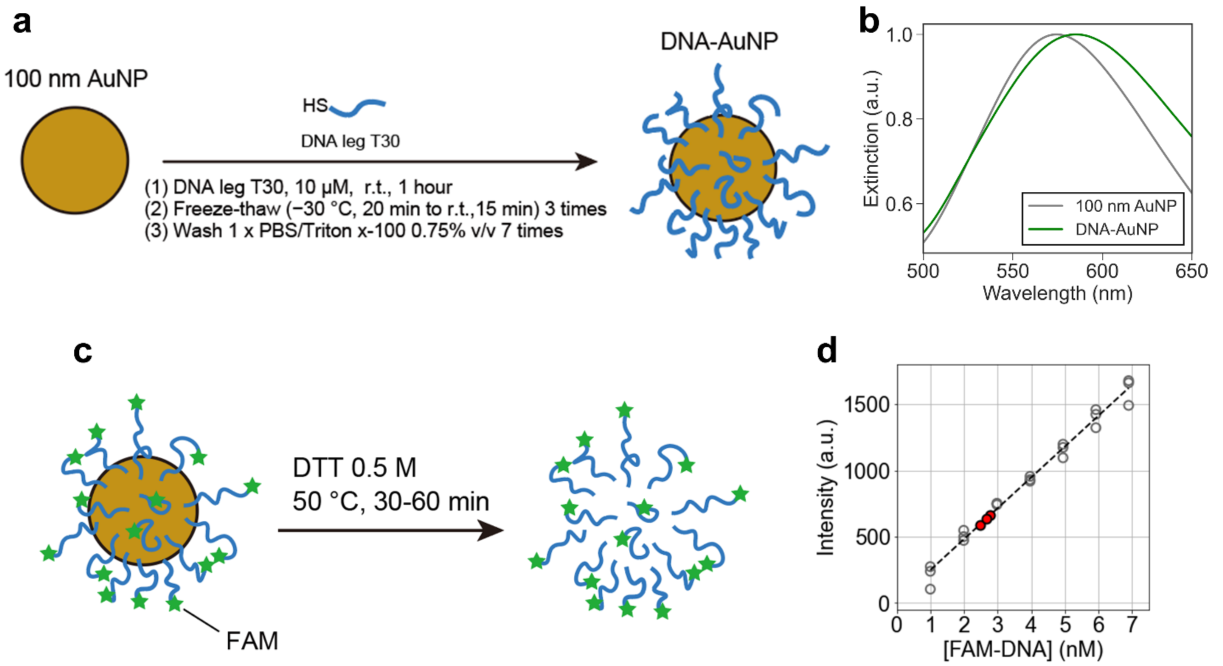


**Supplementary Figure 1. Surface modification of AuNP with DNA and characterization**. **(a)** Schematic illustration of surface modification of AuNP with DNA. The SH group at 5' terminal of DNA binds to the AuNP surface, resulting in oriented DNAs with high surface density. **(b)** Extinction spectra of 100 nm bare AuNP (gray) and DNA-AuNP (green). A slight red shift was observed in DNA-AuNPs. **(c)** Schematic of DNA release reaction by adding 0.5 M DTT. DNA used in this experiment was labeled by fluorescein (FAM) at 3' terminal. **(d)** A calibration curve of [FAM-DNA] and fluorescence intensity at 520 nm ($R^2$ = 0.98). Red colored points show the intensities from diluted solutions of released FAM-DNA and the derived [FAM-DNA]s. The number of DNAs on single AuNP was calculated to be 3500 ± 500 by dividing [FAM-DNA] by the AuNP concentration used in the DNA release experiment. Then, the number was translated into the surface density of 0.11 ± 0.02 molecules nm$^{-2}$ (110000 ± 20000 molecules μm$^{-2}$, N = 3), considering the surface area of a particle with a diameter of 100 nm.

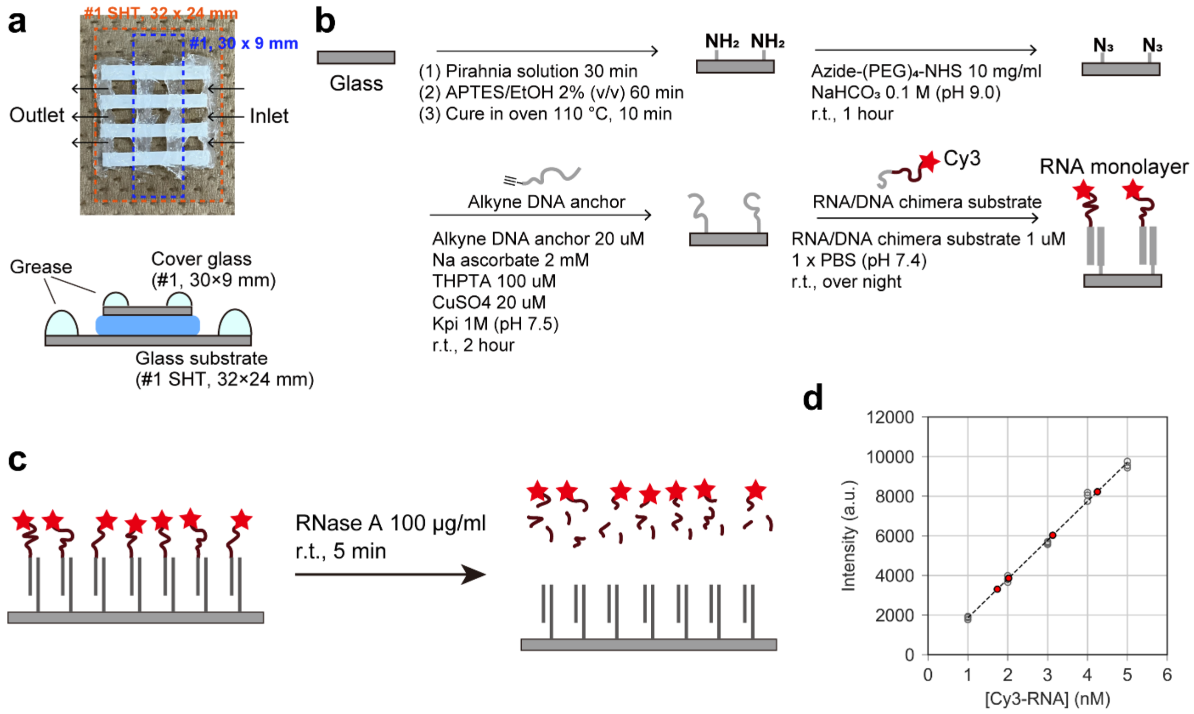


**Supplementary Figure 2. Modification of glass substrate with RNA and characterization. (a)** Photograph of the flow cell used in this study and the illustration of the cross-section. The bottom glass substrate and the top cover glass were separated by double-sided tape with a thickness of 0.62 mm. The bottom glass substrate was modified with RNA. The inlet and outlet of each flow cell were surrounded by grease. **(b)** Scheme of glass substrate modification with RNA . The flow cell was prepared after the APTES treatment. **(c)** Schematic of Cy3-RNA release reaction by adding RNase A, which recognizes and cleaves ssRNAs on the glass substrate. **(d)** A calibration curve of [Cy3-RNA] and fluorescence intensity at 560 nm ($R^2 = 0.997$). Red dot points show the intensities from the solutions with released Cy3-RNA and the derived [Cy3-RNA]s. The total number of Cy3-RNA molecules was translated into the surface density of 0.022 ± 0.008 molecules nm$^{-2}$ (22000 ± 8000 molecules μm$^{-2}$, N = 4) by diving the surface area of the flow cell (2.5 ×9.0 mm$^2$).

**a**

| Condition | [RNase H] (nM) | Cooling fan | Recording rate (frames per second) | Laser power density (nW/μm$^2$) | Localization precision | | |
|---|---|---|---|---|---|---|---|
| | | | | | X-axis (nm) | Y-axis (nm) | $N_{particle}$ |
| 1 | 36 | ON | 20 | 0.57 | 3.1 ± 0.3 | 3.1 ± 0.2 | 26 |
| 2 | 144, 360 | OFF | 20 | 0.57 | 1.0 ± 0.3 | 1.0 ± 0.3 | 24 |
| 3 | 720, 1440 | OFF | 250 | 7.2 | 0.9 ± 0.3 | 0.8 ± 0.3 | 40 |
| 4 | 3600, 7200 | OFF | 1000 | 29 | 0.9 ± 0.3 | 0.9 ± 0.3 | 42 |

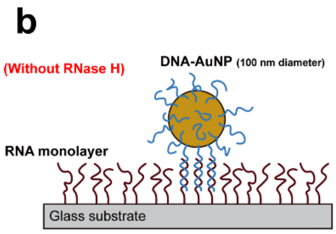


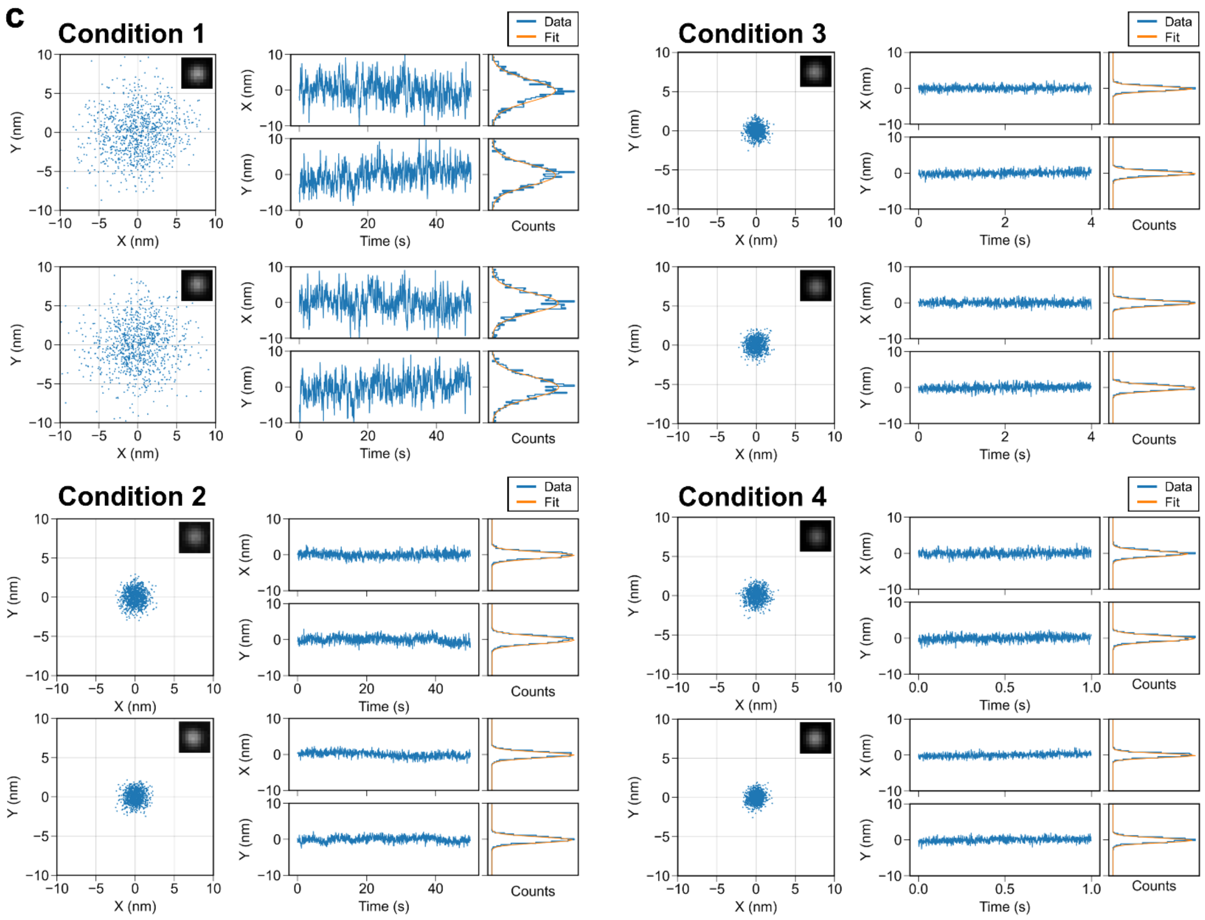


**Supplementary Figure 3. Data recording conditions and localization precisions. (a)** Summary of data recording conditions for different [RNase H]s. **(b)** Schematic illustration of characterization of the localization precision of DNA-AuNP on RNA monolayer. DNA-AuNPs were attached to the RNA monolayer, and their positions were measured without the addition of RNase H. **(c)** Left: Examples of centroid trajectory of DNA-AuNP (Inset: A scattering images of the DNA-AuNP). Middle: Time courses of X- and Y-coordinates. Right: Histograms of X- and Y-coordinates. Localization precisions were calculated as the standard deviations of the fitted Gaussian functions (orange lines).

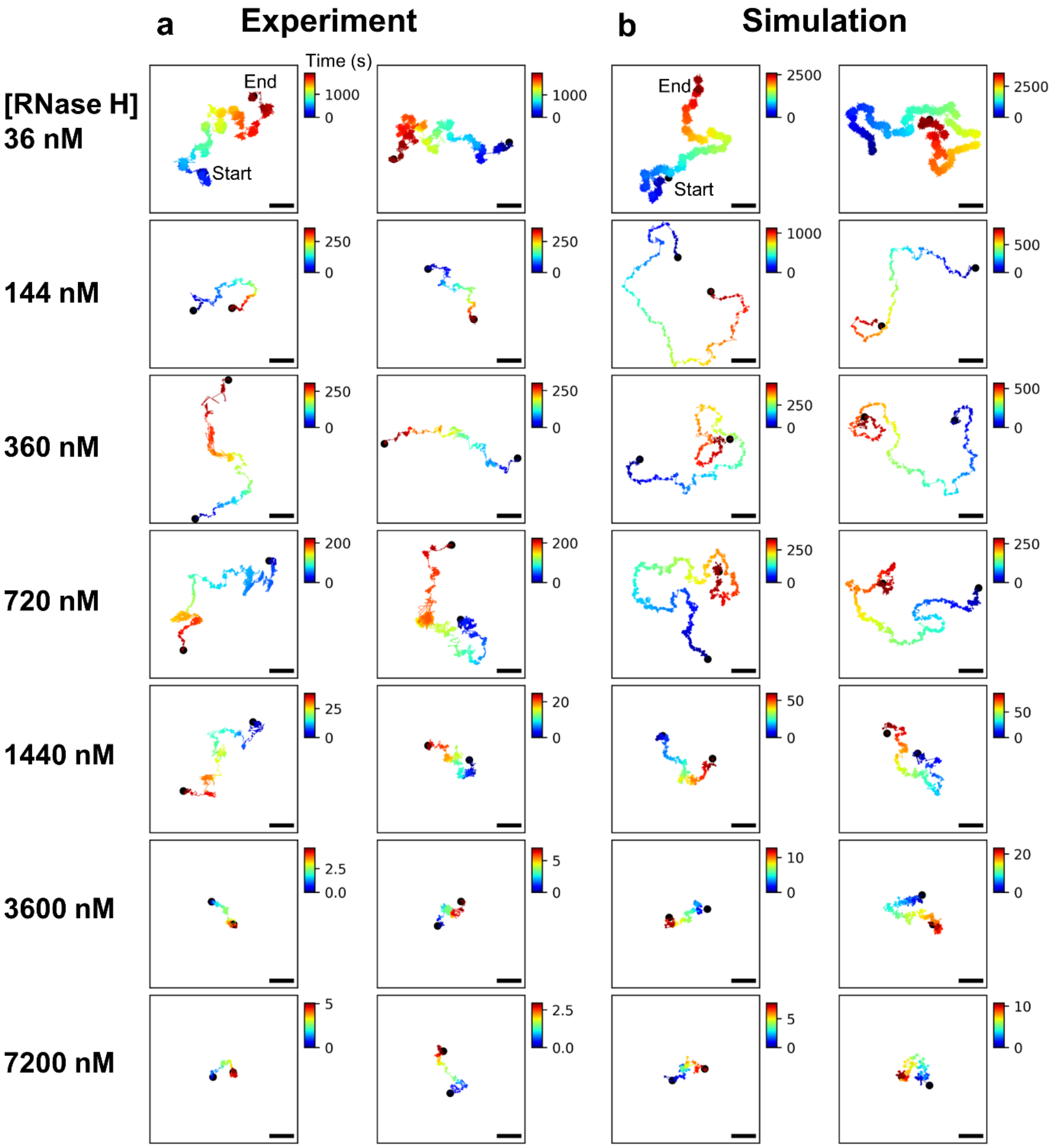


**Supplementary Figure 4. Typical trajectories of experiment and simulation at each [RNase H].** Examples of experimental (**a**) and simulation (**b**) trajectories. Two trajectories are shown for each [RNase H] condition in the experiments and simulations. Scalebar: 100 nm. Gaussian noise with standard deviation ($\sigma_{\text{noise}}$) of 6 nm was added to the simulation trajectories at 36 nM RNase H, and that with $\sigma_{\text{noise}}$ of 2 nm was added to the simulation trajectories at 144, 360, 720, 1440, 3600, and 7200 nM RNase H (see Supplementary Fig. 9 and 10 for details).

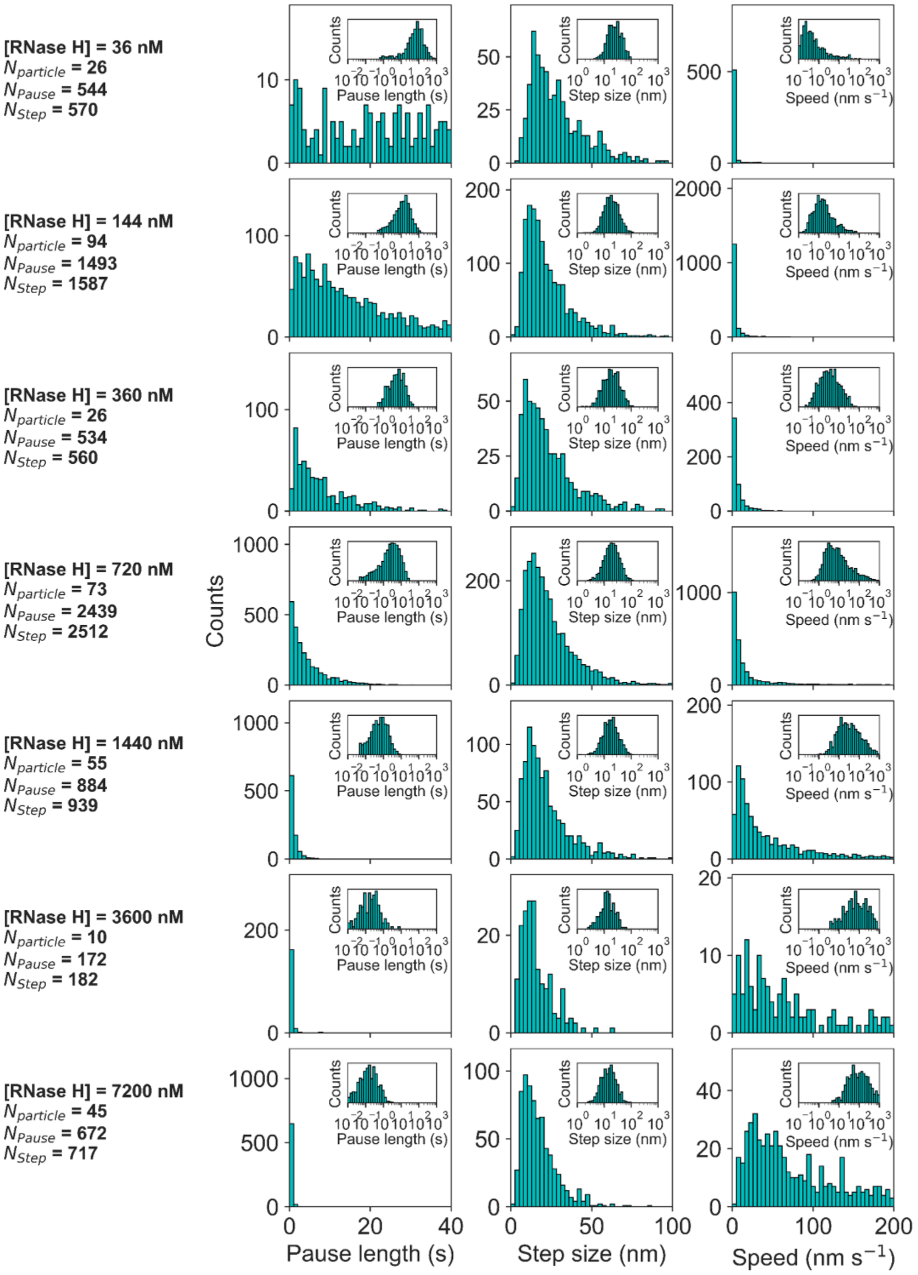


**Supplementary Figure 5. Distributions of pause length, step size, and speed at each [RNase H] for the experimental data.** Distributions in log scale are also shown in the insets.

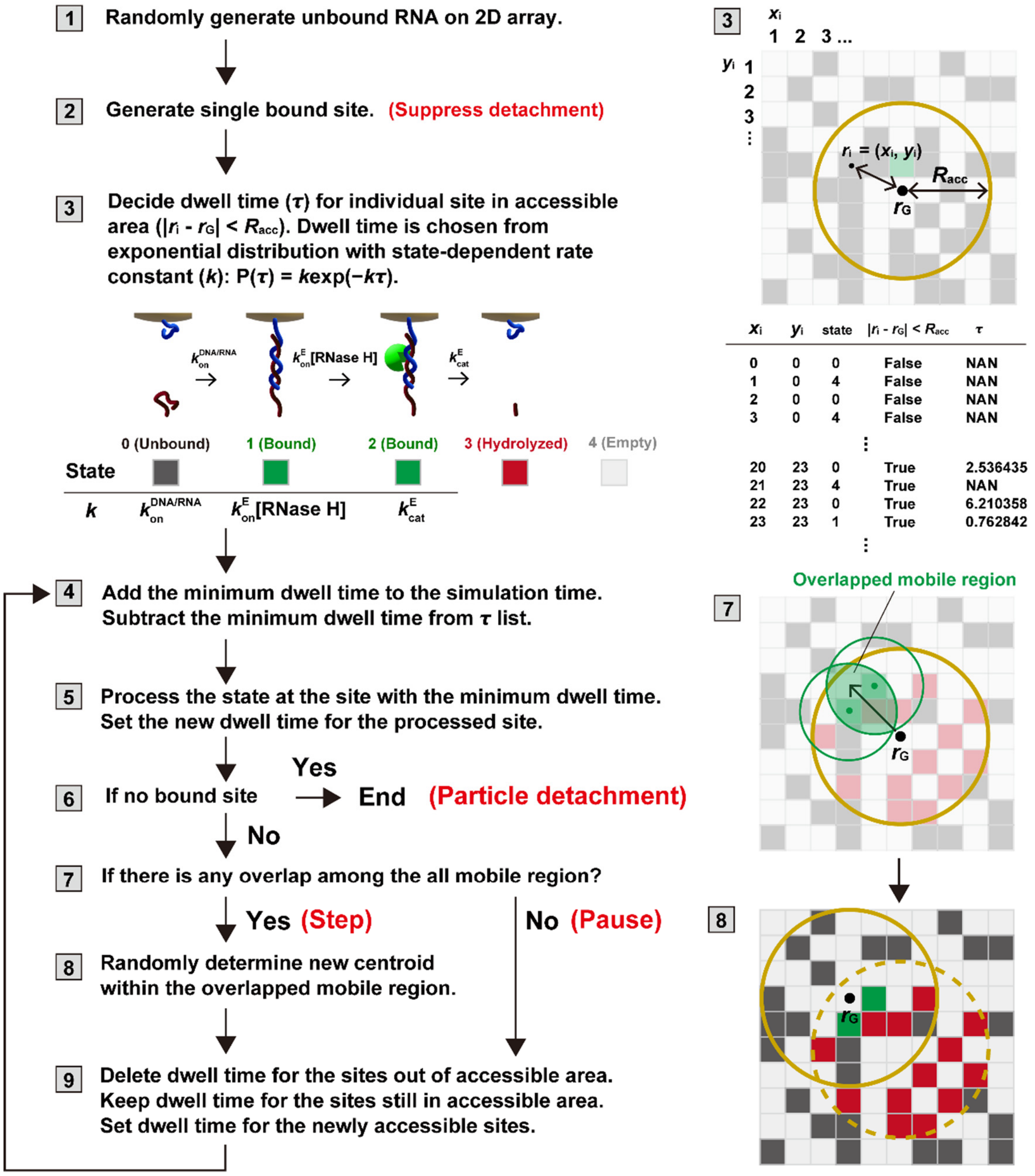


**Supplementary Figure 6. Flowchart of geometry-based kinetic simulation.** Flow chart and schematic illustration are shown. The simulation framework was implemented in Python 3. The NumPy package was used for operations on two-dimensional state distribution, and the Pandas package was used to summarize the time course of X- and Y-coordinates at each step. To obtain a dwell time from an exponential distribution P(t) = $k$exp(-$k\tau$), where $k$ is the state-dependent rate constant, the expovariate() function provided by the random package was utilized. The matplotlib package was employed for plotting the results.

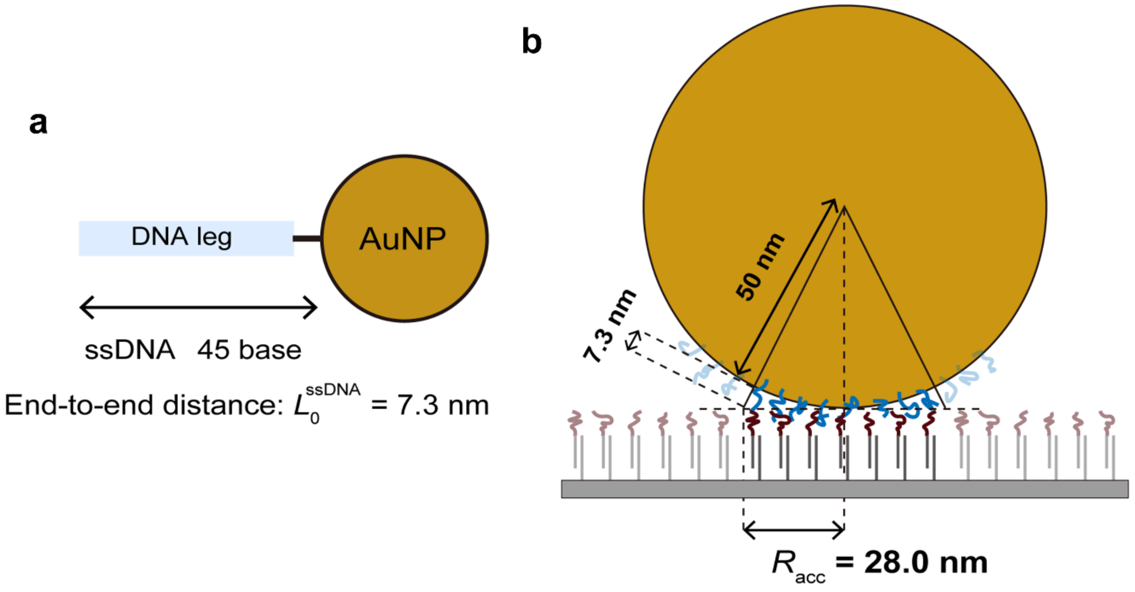


**Supplementary Figure 7. Geometric model to calculate the accessible radius. (a)** Schematic illustration of the end-to-end distance of single-stranded DNA ($L_0^{ssDNA}$) with 45 bases. **(b)** Schematic of the geometric model to calculate the accessible radius ($R_{acc}$) used in the simulation.

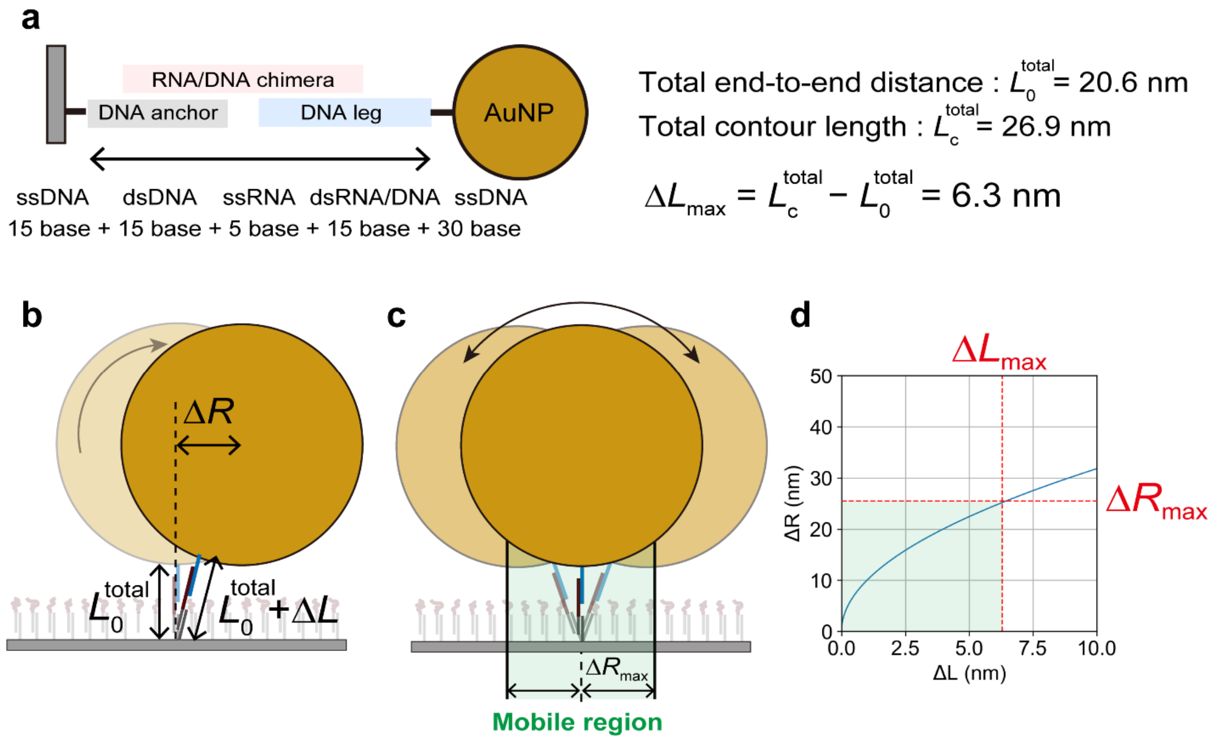


**Supplementary Figure 8. Configuration of the mobile region for the motion of DNA-AuNP motor. (a)** Schematic illustration of the DNA/RNA duplex to estimate the end-to-end distance and the contour length. Calculated total end-to-end distance ($L_0^{total}$) without applied force and total contour length ($L_c^{total}$) were 20.6 nm and 26.9 nm, respectively. **(b)** Geometric model used to calculate the mobile region for DNA/RNA duplex at the bound site in the simulation. We assumed that the AuNP moves laterally ($\Delta R$) with rolling by thermal agitation, which accompanies the increase ($\Delta L$) of the end-to-end distance. **(c)** We assumed that the end-to-end distance varies in the range between $L_0^{total}$ and $L_c^{total}$, where the maximum $\Delta L$ ($\Delta L_{max}$) is 6.3 nm (26.9 – 20.6 nm). **(d)** Relationship between $\Delta L$ and $\Delta R$. A green-shadowed region corresponds to the mobile region of a single RNA/DNA bound site. When $\Delta L_{max}$ is 6.3 nm, the maximum $\Delta R$ ($\Delta R_{max}$) becomes 25.2 nm. In our simulation, the motor steps are allowed when the mobile regions for all bound sites overlap.

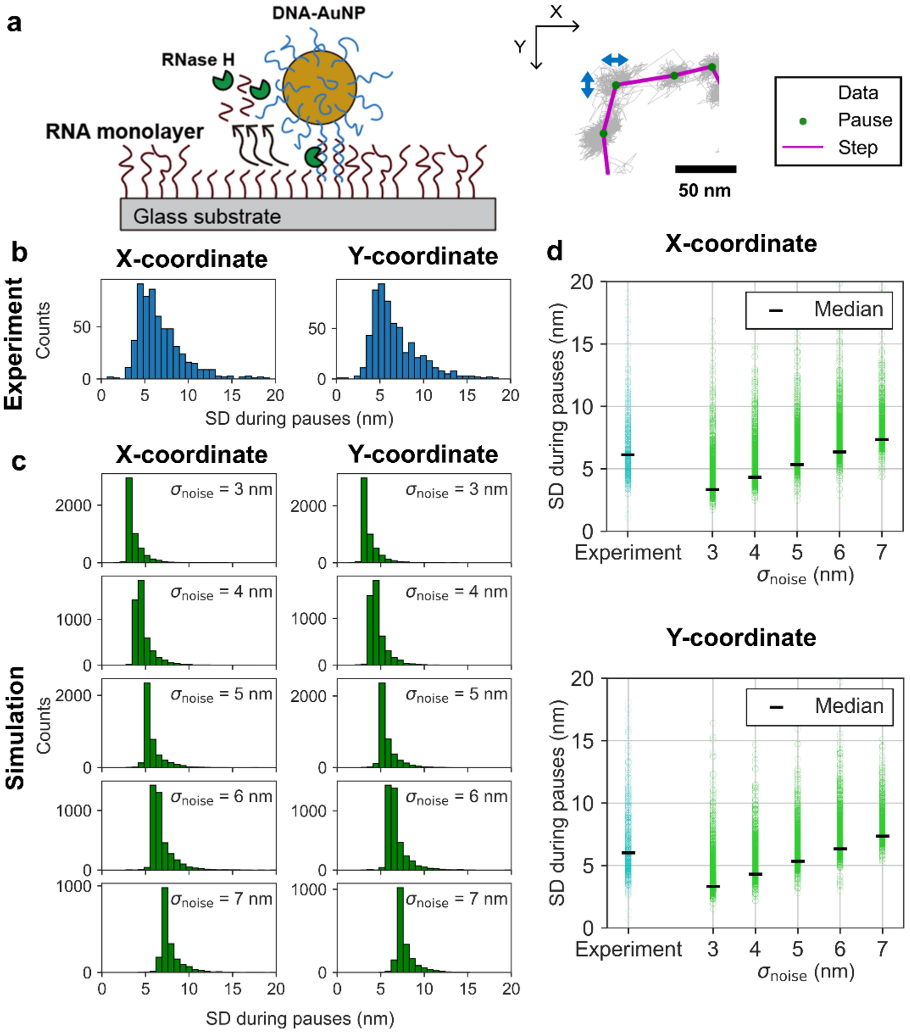


**Supplementary Figure 9. Optimization of noise added to simulation trajectory to reproduce experimental trajectory with camera fan on. (a)** Schematic illustration of active DNA-AuNP motor in the presence of RNase H (left) and a cropped trajectory (gray) superimposed with steps (magenta lines) and pauses (green dots) detected by the step-finding algorithm (right). [RNase H]: 36 nM. Recording rate: 20 frames per second (fps). **(b)** Histograms of standard deviation (SD) of X- and Y-coordinates during pauses of experimental trajectories. **(c)** Histograms of SD of X- and Y-coordinates during pauses of simulation trajectories with Gaussian noise ($\sigma_{noise}$ of 3, 4, 5, 6, or 7 nm). **(d)** Summarized plots of the SD during pauses in the experiment (cyan circles) and simulations (green circles). Median values are calculated for each condition and shown by black horizontal bars. Median values for X- and Y-coordinates in the experiment were calculated to be 6.1 and 6.0 nm, respectively. Optimal $\sigma_{noise}$ was determined to be 6 nm with a median value of 6.4 nm for both X- and Y-coordinates.

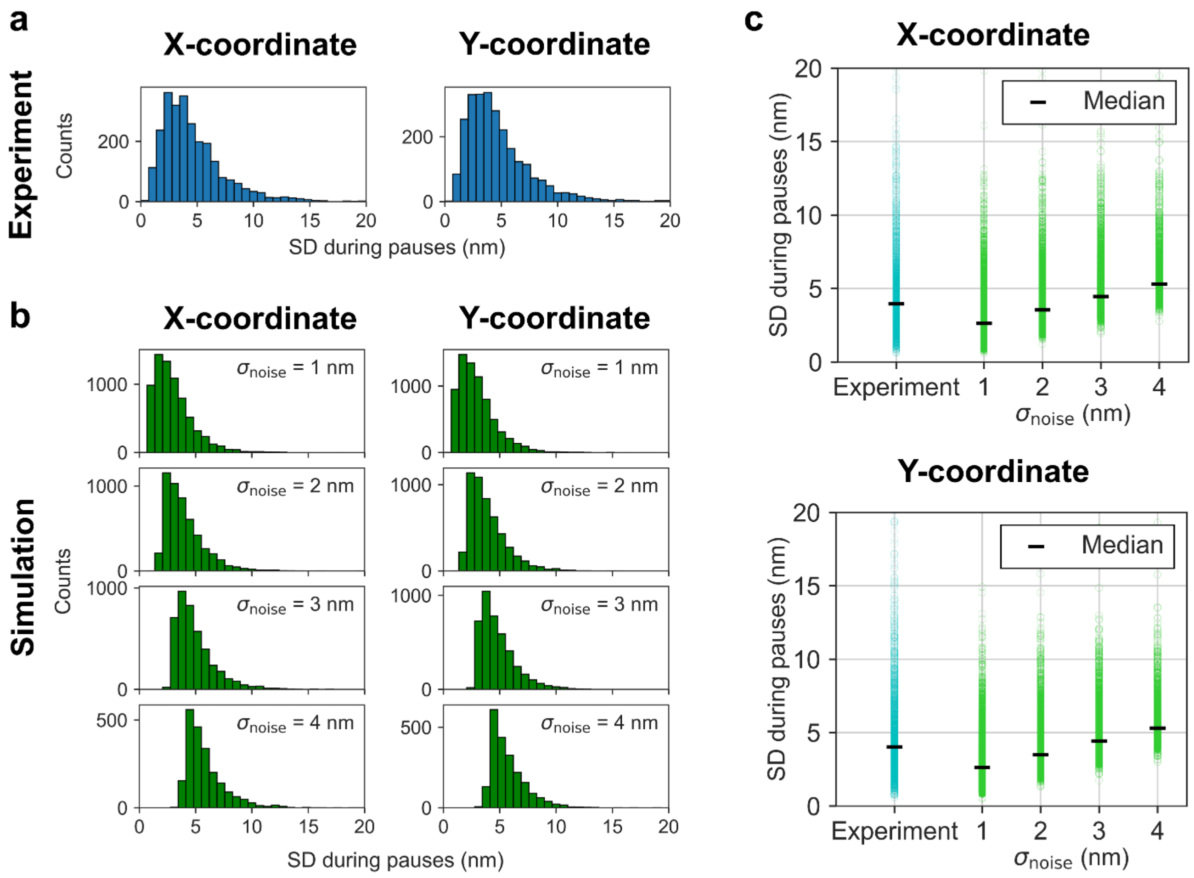


**Supplementary Figure 10. Optimization of noise added to simulation trajectory to reproduce experimental trajectory with camera fan off. (a)** Histograms of standard deviation (SD) of X- and Y-coordinates during pauses of experimental trajectories. **(b)** Histograms of SD of X- and Y-coordinates during pauses of simulation trajectories with Gaussian noise ($\sigma_{noise}$ of 1, 2, 3, or 4 nm). **(d)** Summarized plots of the SD during pauses in the experiment (cyan circles) and simulations (green circles). Median values are calculated for each condition and shown by black horizontal bars. Median values for X- and Y-coordinates in the experiment were calculated to be 4.0 nm for both X- and Y-coordinates. Optimal $\sigma_{noise}$ was determined to be 2 nm with median values of 3.6 and 3.5 nm for X- and Y-coordinates, respectively.

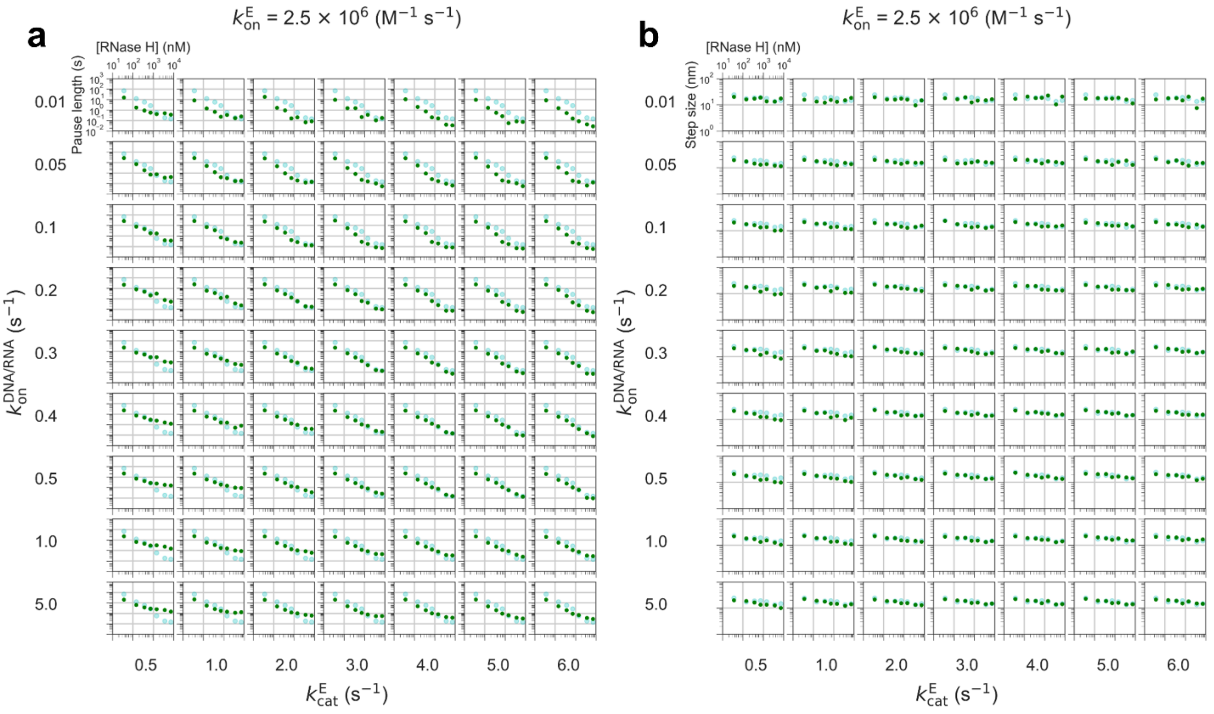


**Supplementary Figure 11. [RNase H] dependence of pause length (a) and step size (b) in kinetic simulations with $k_{on}^{E} = 2.5 \times 10^{6}$ M$^{-1}$s$^{-1}$.** Different sets of $k_{on}^{DNA/RNA}$ and $k_{cat}^{E}$ were used. Green and light blue plots show the simulation and experimental data, respectively.

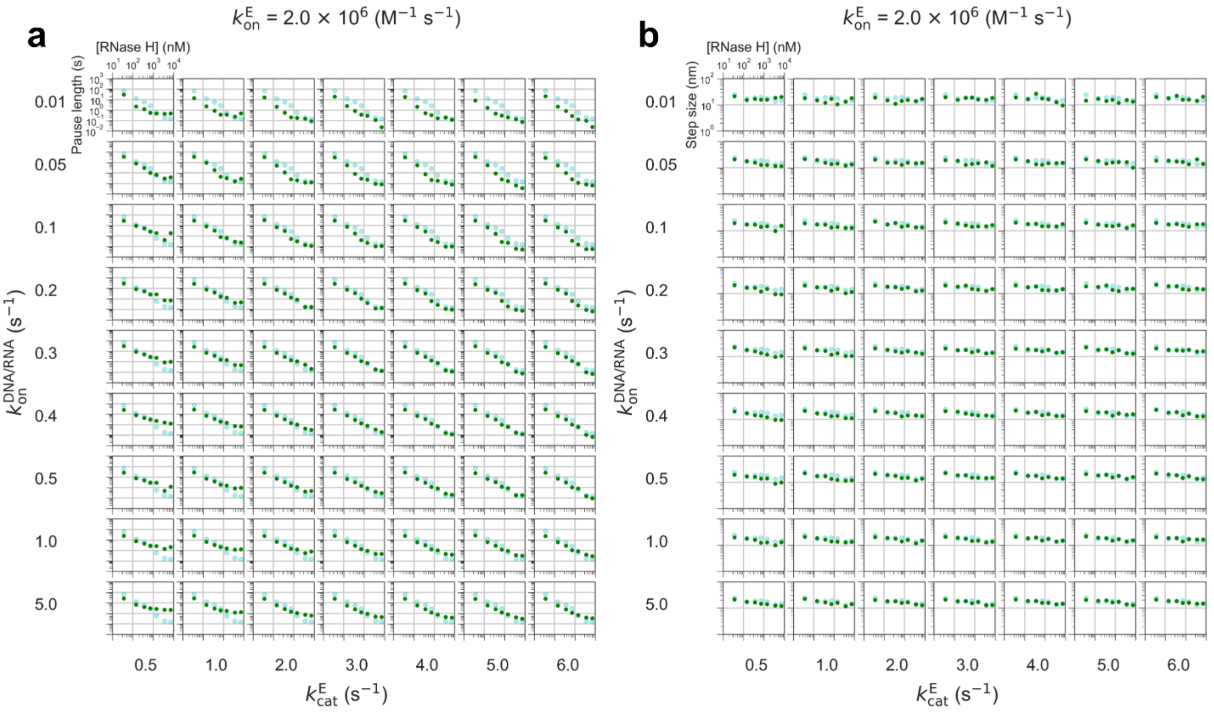


**Supplementary Figure 12. [RNase H] dependence of pause length (a) and step size (b) in kinetic simulations with $k_{on}^{E} = 2.0 \times 10^{6}$ M$^{-1}$s$^{-1}$.** Different sets of $k_{on}^{DNA/RNA}$ and $k_{cat}^{E}$ were used. Green and light blue plots show the simulation and experimental data, respectively.

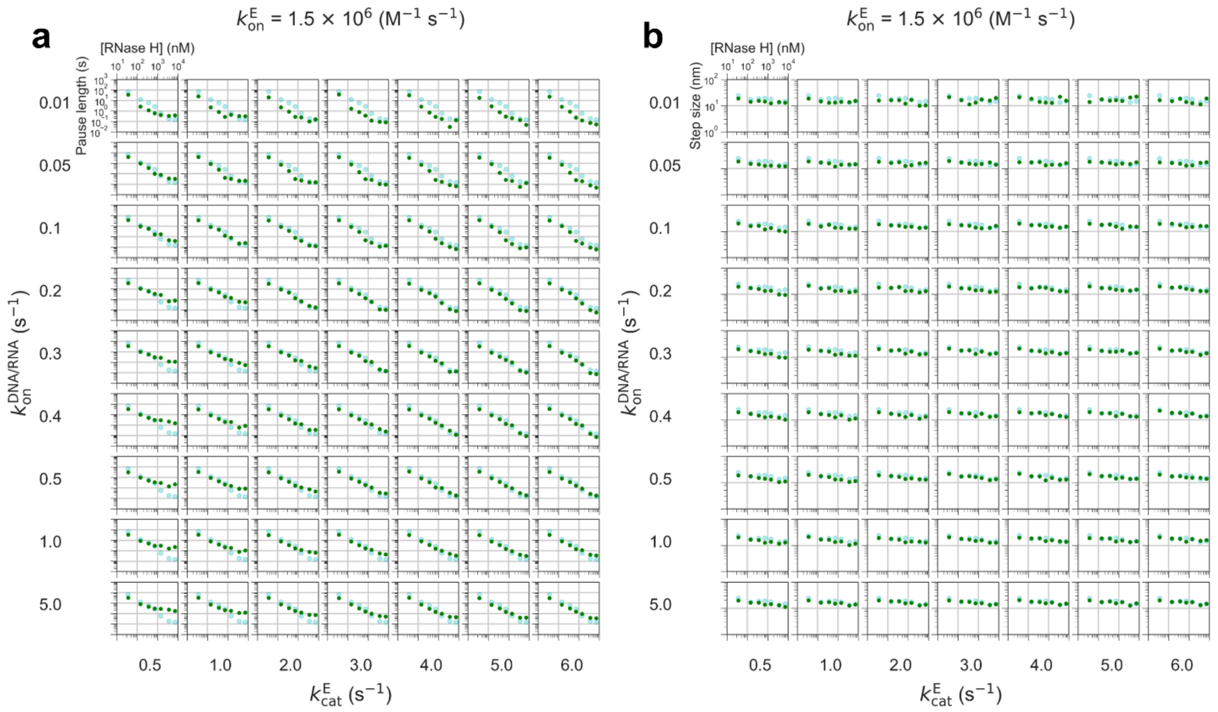


**Supplementary Figure 13. [RNase H] dependence of pause length (a) and step size (b) in kinetic simulations with $k_{on}^{E}$ = 1.5 × $10^6$ $M^{-1}s^{-1}$.** Different sets of $k_{on}^{DNA/RNA}$ and $k_{cat}^{E}$ were used. Green and light blue plots show the simulation and experimental data, respectively.

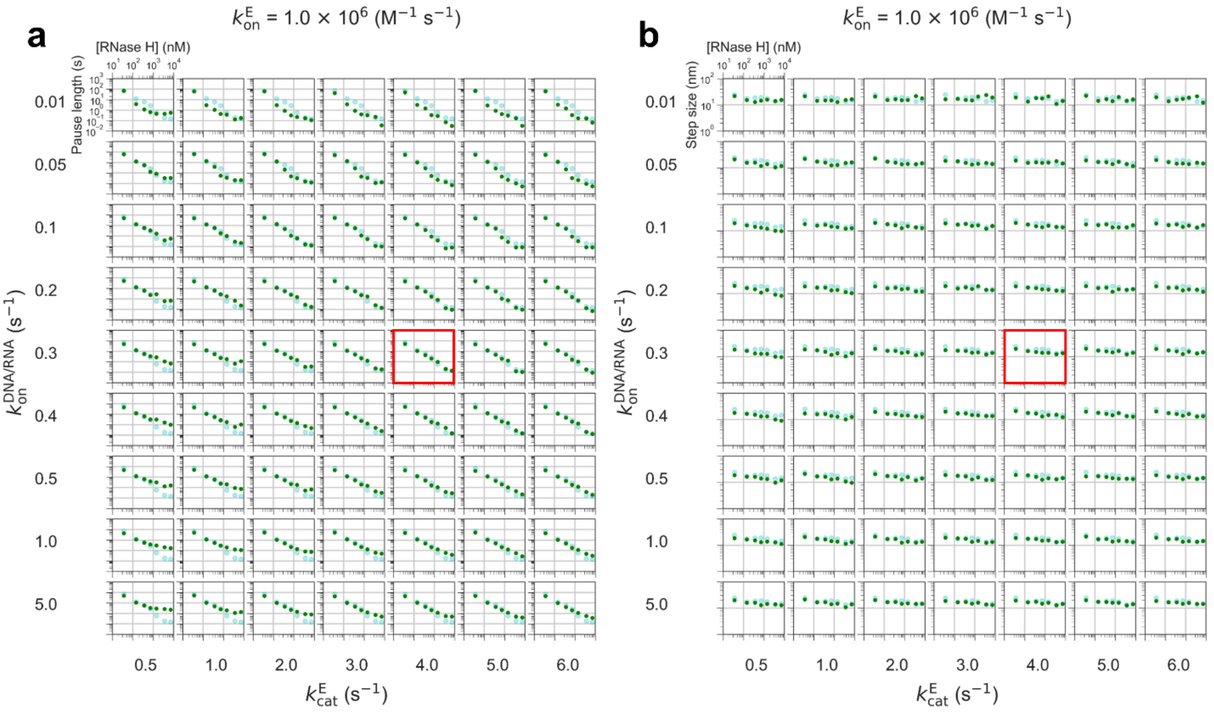


**Supplementary Figure 14. [RNase H] dependence of pause length (a) and step size (b) in kinetic simulations with $k_{on}^{E}$ = 1.0 × 10$^{6}$ M$^{-1}$s$^{-1}$.** Different sets of $k_{on}^{DNA/RNA}$ and $k_{cat}^{E}$ were used. Green and light blue plots show the simulation and experimental data, respectively. Results of a simulation with $k_{on}^{DNA/RNA}$ = 0.3 s$^{-1}$, $k_{on}^{E}$ = 1.0×10$^{6}$ M$^{-1}$ s$^{-1}$, and $k_{cat}^{E}$ = 4 s$^{-1}$, which showed the minimum mean squared error (MSE) against experimental data, are highlighted by red boxes.

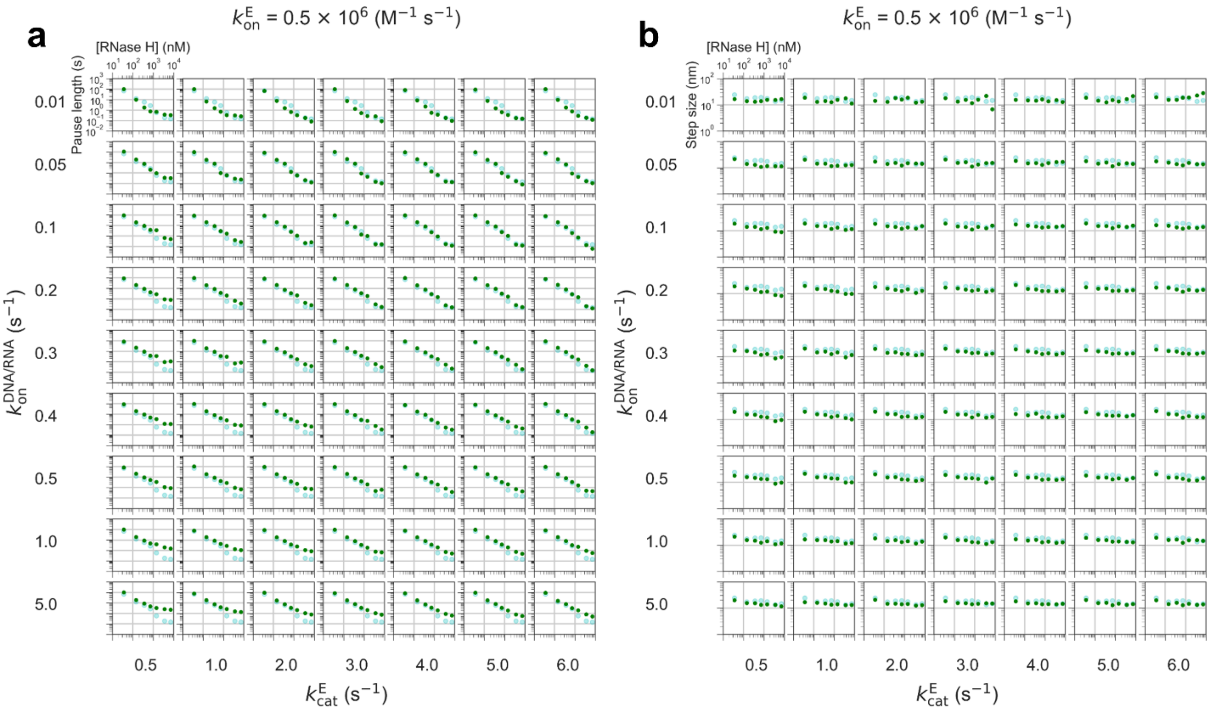


**Supplementary Figure 15. [RNase H] dependence of pause length (a) and step size (b) in kinetic simulations with $k_{on}^{E}$ = 0.5 × 10$^{6}$ M$^{-1}$s$^{-1}$.** Different sets of $k_{on}^{DNA/RNA}$ and $k_{cat}^{E}$ were used. Green and light blue plots show the simulation and experimental data, respectively.

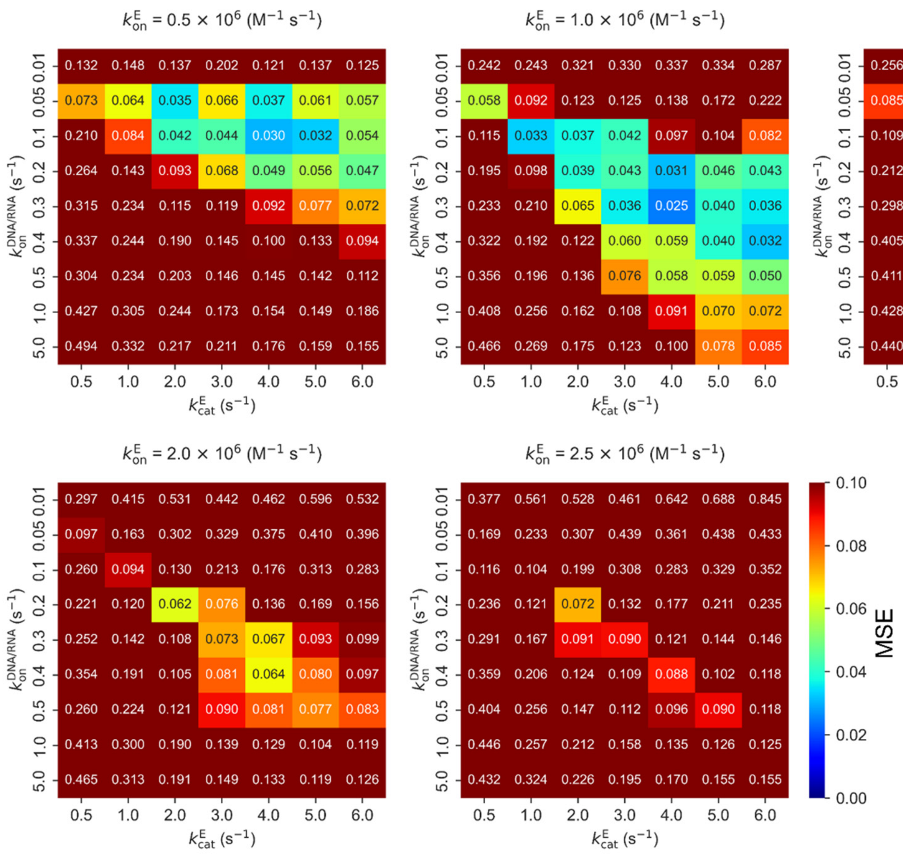


**Supplementary Figure 16. Optimization of rate constants by simulation-based fitting.** Mean squared error (MSE) of the pause length and the step size between the simulation and the experiment were evaluated to determine the optimal set of the rate constants. The optimal set which showed the MSE of 0.025 (highlighted by blue box) was $k_{on}^{DNA/RNA} = 0.3\ s^{-1}$, $k_{on}^{E} = 1.0\times10^{6}\ M^{-1}\ s^{-1}$, and $k_{cat}^{E} = 4\ s^{-1}$.

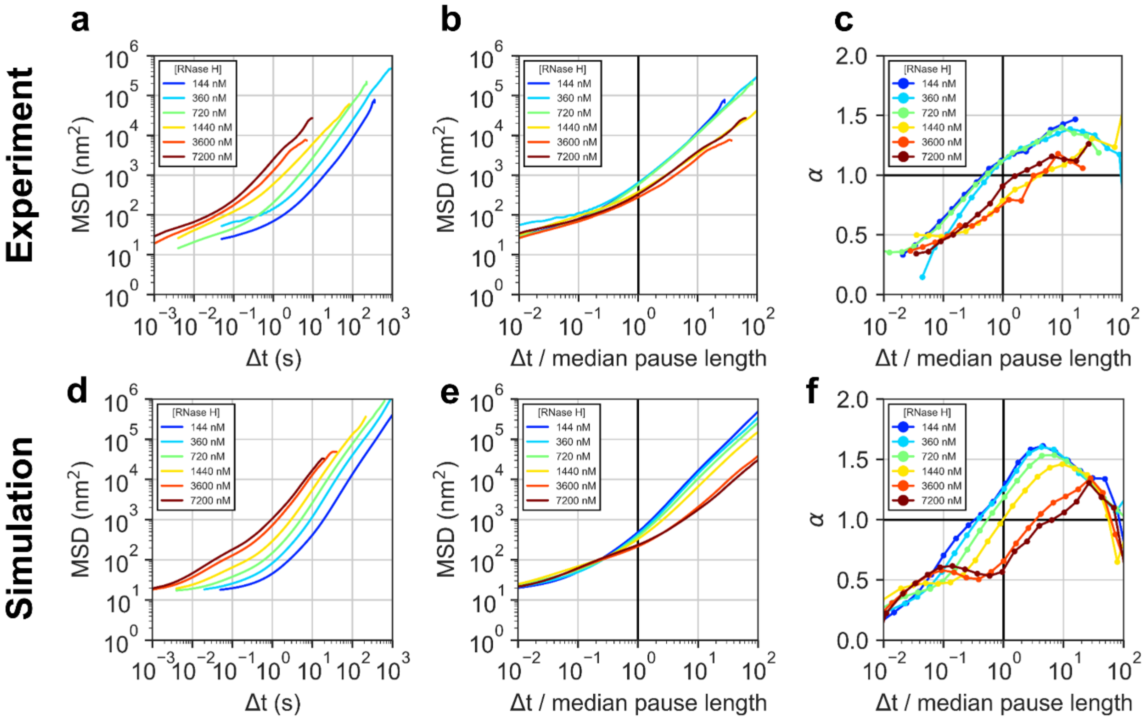


**Supplementary Figure 17. MSD analysis.** (**a, d**) Log-log plots of the mean square displacement (MSD) as a function of time increment ($\Delta t$) for experimental (a) and simulation (d) data at each [RNase H]. (**b, e**) Log-log plots of MSD vs. normalized $\Delta t$ ($\Delta t$ divided by median pause lengths calculated in Fig. 2) for experimental (b) and simulation (e) data at each [RNase H]. (**c, f**) Plots of the anomalous factor α vs. normalized $\Delta t$ for experimental (c) and simulation (f) data at each [RNase H].

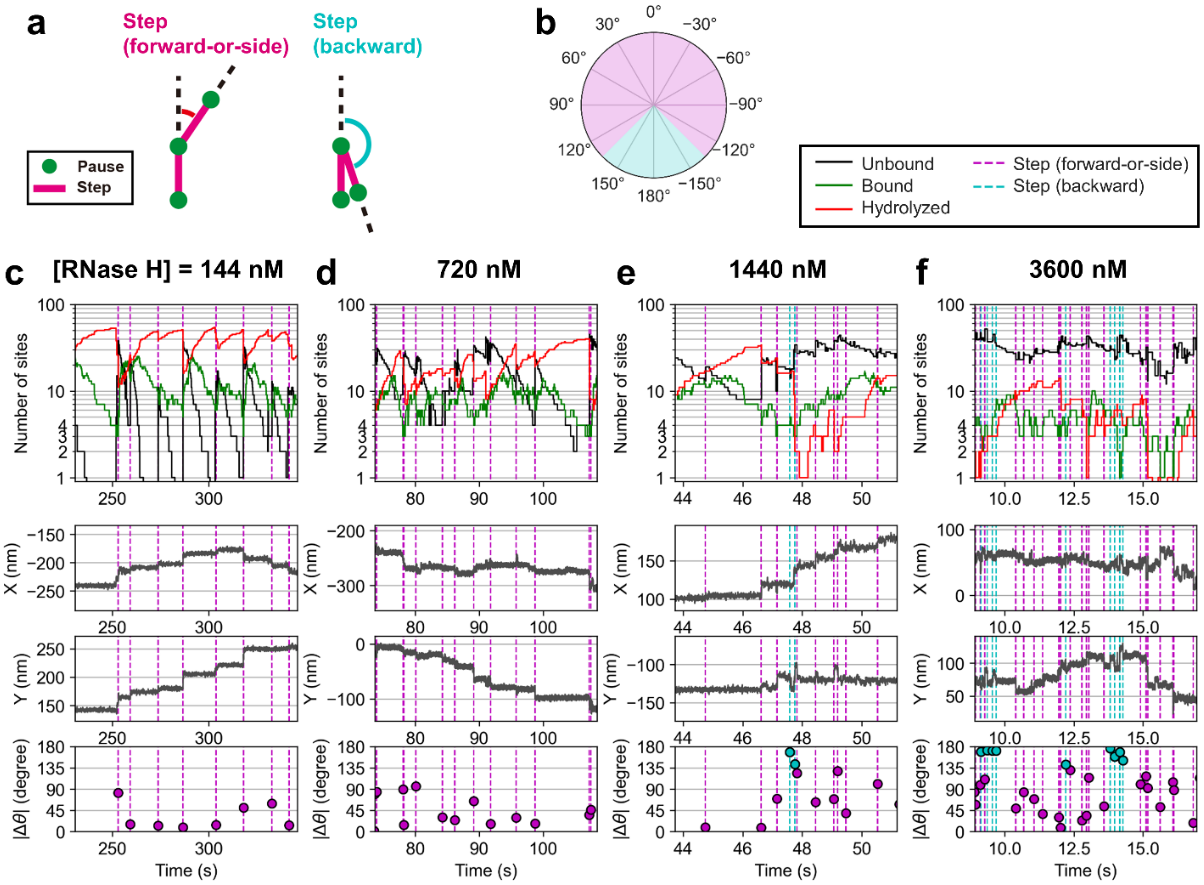


**Supplementary Figure 18. [RNase H] dependence of time-course of the number of RNA sites, X- and Y-coordinates, and step angles (Δ*θ*) in the simulation.** (**a**) Schematic illustration of step angle analysis. **(b)** A polar plot showing the angle regions of forward-or-side step (pink shaded) and backward steps (cyan shaded). Steps with |Δ*θ*| < 145° and > 145° were classified as forward-or-side and backward steps, respectively. (**c**‒**f**) Examples of time-course of the number of RNA sites, X- and Y-coordinates, and |Δ*θ*| at 144 (c), 720 (d), 1440 (e), and 3600 (f) nM RNase H. Forward-or-side and backward steps are shown as magenta and cyan dotted lines, respectively.

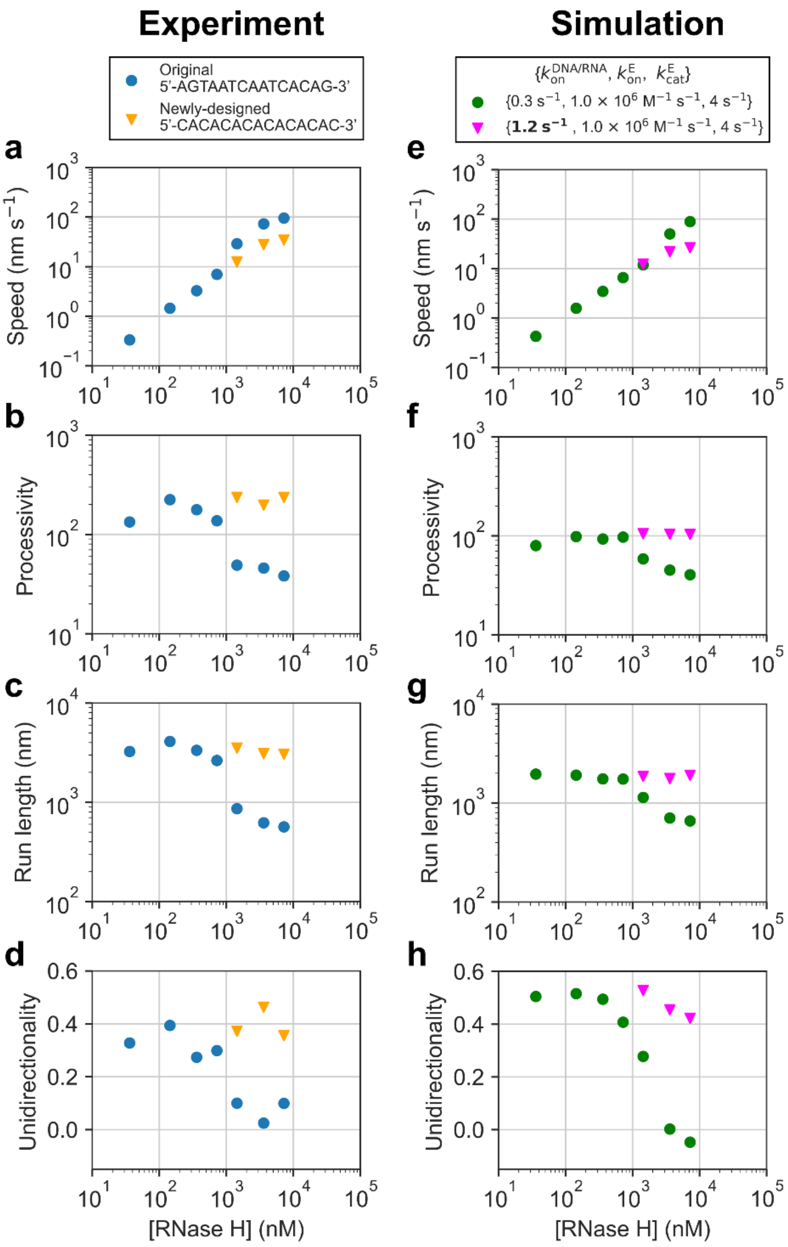


**Supplementary Figure 19. Summary of [RNase H] dependence of motor performance. (a-d)** [RNase H] dependence of speed (a), processivity (b), run-length (c), and unidirectionality (d) in the experiments. Blue circles and orange triangles correspond to the data using original and newly-designed DNA/RNA sequence, respectively. **(e-h)** [RNase H] dependence of speed (e), processivity (f), run-length (g), and unidirectionality (h) in the simulation. Green circles correspond to the data using $k_{on}^{DNA/RNA} = 0.3\ s^{-1}$, $k_{on}^{E} = 1.0 \times 10^{6}\ M^{-1}s^{-1}$, and $k_{cat}^{E} = 4\ s^{-1}$. Pink triangles correspond to the data using $k_{on}^{DNA/RNA} = 1.2\ s^{-1}$, $k_{on}^{E} = 1.0{\times}10^{6}\ M^{-1}s^{-1}$, and $k_{cat}^{E} = 4\ s^{-1}$.

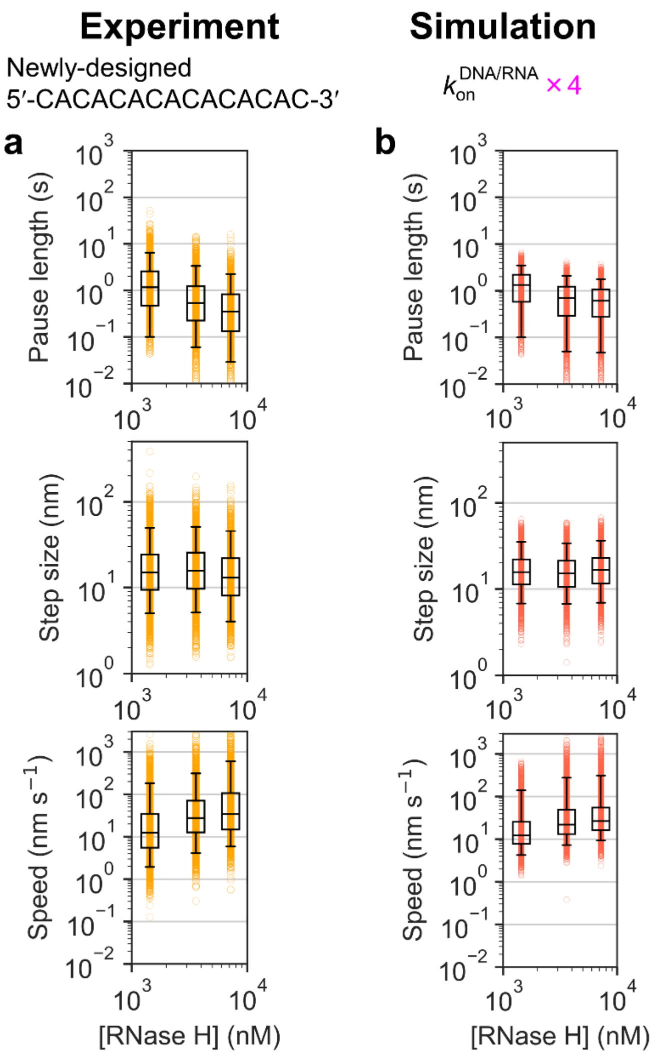


**Supplementary Figure 20. Step analysis of the DNA-AuNP motor with large hybridization rates. (a)** [RNase H] dependence of pause length (top), step size (middle), and speed (bottom) in the experiment at 1440, 3600, and 7200 nM RNase H in the experiments using newly-designed DNA/RNA sequence. All the data are shown by cyan circles. Box plot shows interquartile range (IQR). Error bars show 5$^{th}$/95$^{th}$ percentile for each [RNase H] condition. Recording rates were 250, 1000, and 1000 frames per second (fps) for 1440, 3600, and 7200 nM RNase H, respectively. **(b)** [RNase H] dependence of pause length (top), step size (middle), and speed (bottom) in the simulation using $k_{on}^{DNA/RNA} = 1.2\ s^{-1}$, $k_{on}^{E} = 1.0 \times 10^6\ M^{-1}s^{-1}$, and $k_{cat}^{E} = 4\ s^{-1}$ where $k_{on}^{DNA/RNA}$ is 4-times larger than that used in Fig. 2.

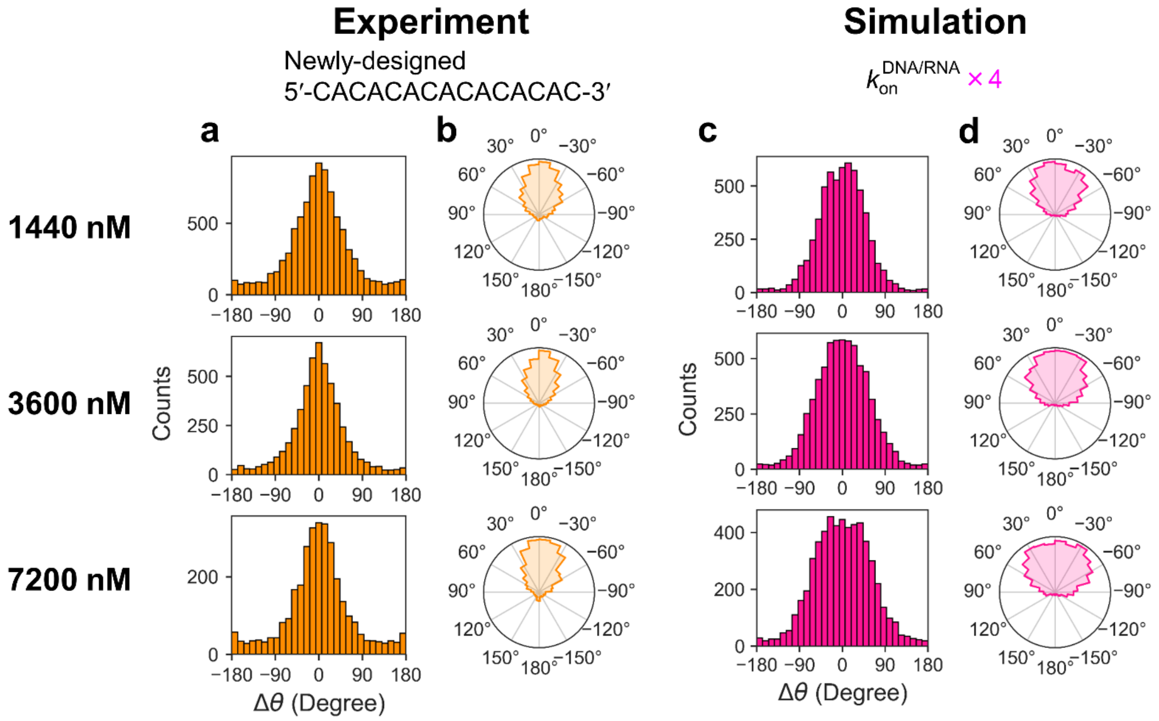


**Supplementary Figure 21. Step angle analysis of the DNA-AuNP motor with large hybridization rates. (a, b)** Step angle histograms and polar plots at 1440, 3600, and 7200 nM RNase H in the experiment using newly-designed DNA/RNA sequence. **(c, d)** Step angle histograms and polar plots in the simulation using ($k_{on}^{DNA/RNA} = 1.2$ s$^{-1}$, $k_{on}^{E} = 1.0\times10^{6}$ M$^{-1}$s$^{-1}$, and $k_{cat}^{E} = 4$ s$^{-1}$ where $k_{on}^{DNA/RNA}$ is 4-times larger than that used in Fig. 4.

**Supplementary References**


1. Roth, E., Glick Azaria, A., Girshevitz, O., Bitler, A. & Garini, Y. Measuring the Conformation and Persistence Length of Single-Stranded DNA Using a DNA Origami Structure. *Nano Lett.* **18**, 6703-6709, (2018).
2. Zhang, C. *et al.* The Mechanical Properties of RNA-DNA Hybrid Duplex Stretched by Magnetic Tweezers. *Biophys. J.* **116**, 196-204, (2019).
3. Chen, H. *et al.* Ionic strength-dependent persistence lengths of single-stranded RNA and DNA. *Proc. Natl. Acad. Sci. USA* **109**, 799-804, (2012).
4. Howard, J. *Mechanics of motor proteins and the cytoskeleton* (Sinauer Associates, 2001).
5. Metzler, R., Jeon, J. H., Cherstvy, A. G. & Barkai, E. Anomalous diffusion models and their properties: non-stationarity, non-ergodicity, and ageing at the centenary of single particle tracking. *Phys. Chem. Chem. Phys.* **16**, 24128-24164, (2014).
6. Korosec, C. S. *et al.* Substrate stiffness tunes the dynamics of polyvalent rolling motors. *Soft Matter* **17**, 1468-1479, (2021).
7. Bazrafshan, A. *et al.* DNA Gold Nanoparticle Motors Demonstrate Processive Motion with Bursts of Speed Up to 50 nm Per Second. *ACS Nano* **15**, 8427-8438, (2021).
8. Yehl, K. *et al.* High-speed DNA-based rolling motors powered by RNase H. *Nat. Nanotechnol.* **11**, 184-190, (2016).
